# Hallmarks of adaptive immunity in a metastatic clear cell renal cell carcinoma (ccRCC) long-term elite survivor

**DOI:** 10.64898/2025.12.19.695541

**Authors:** Felicia Tucci, Edward Drydale, James Bancroft, Sakina Amin, Jane Pernes, Giulia Gardumi, Tinashe Kanyowa, Maria Lima, Nadia Nasreddin, Olivia Lombardi, Simon Leedham, Aniko Rendek, Lisa Browning, Emmanuel Bugarin Estrada, Ruddy Montandon, Santiago Revale, Hubert Slawinski, David Mole, Andy Protheroe, Shivan Sivakumar, Rachael Bashford-Rogers

## Abstract

Clear cell renal cell carcinoma (ccRCC) presents with metastatic disease in nearly one-third of patients, with a 5-year survival of only ∼10%. Although anti-angiogenic therapies and immune checkpoint inhibitors (ICIs) have improved outcomes, therapeutic resistance and immune-related adverse events limit durable responses. Exceptional long-term survivors provide a unique opportunity to uncover mechanisms of sustained anti-tumour immunity and guide therapeutic strategies.

Here, we present a spatial and temporal dissection of adaptive immunity in an extraordinary 16-year ccRCC survivor, integrating single-cell sequencing, multiplex immunofluorescence, B and T cell receptor (BCR/TCR) repertoire analysis, and antibody profiling. We reveal persistent and spatially coordinated B and T cell clones over 16 years, indicating long-term immune memory and surveillance throughout tumour evolution.

B cell analysis identified pronounced clonal expansions involving intrinsically autoreactive IGHV4-34 B cells undergoing ongoing germinal centre (GC) maturation in tumour-draining lymph nodes (dLNs) and spleen. These clones progressively lost autoreactivity while acquiring tumour-specificity through somatic hypermutation (SHM), consistent with clonal redemption. Redeemed B cells displayed enhanced antigen-presenting functions and localised to tertiary lymphoid structures (TLS), where they interacted with CD8⁺ T cells, as inferred from doublet analysis. In parallel, persistent CD8^+^ effector T cell clones highlighted the contribution of long-lived lymphocytes to tumour control.

Comparison of the primary tumour and pancreatic metastatic with paired tumour-free dLNs showed that TLS mirrored GC functionality, particularly for PD-1^+^ CD8^+^ T cells, while TLS maturation was stroma-dependent and impaired at the metastatic site. Finally, tumour-derived BCRs revealed antibody binding to tumour specific “public” antigens without cross-reactivity to normal tissue. Together, these findings establish clonal persistence, TLS engagement, and coordinated B-T cell immune surveillance as hallmarks of durable tumour control, providing a framework for next-generation antibody therapeutics and patient stratification in metastatic ccRCC.

**What is already known on this topic.:** Tertiary lymphoid structures within the tumour microenvironment (TME) and clonal expansions of CD8^+^ T cells are associated with better survival and improved response to immunotherapies in ccRCC. However, identifying markers of prolonged survival is challenging due to the rarity of long-term survivors (LTS), and the predominance of studies focused on T cell immunity, leaving B cell contribution almost unexplored.

**What this study adds.:** This study provides an integrated analysis of B and T cell responses in an exceptional 16-year LTS with metastatic ccRCC, revealing key immunologic features of sustained anti-tumour immunity. Beyond T cell dynamics, it highlights the multifaceted role of B cells as antigen presenting cells, antibody producers, and modulators of CD8^+^ T cell effector functions through direct interactions within TLS. Importantly, it reveals potential tumour-specific antigens driving humoral responses.

**How this study might affect research, practice or policy.:** By uncovering mechanisms of sustained tumour control, this work establishes a framework for identifying adaptive immune biomarkers of long-term survival and refining patient stratification in metastatic ccRCC. Multimodal assessment of B cell specificity and spatial interactions between re-educated B cells and CD8^+^ T cells within TLS introduces a new paradigm for understanding coordinated B and T cell responses. These insights have direct implications for antibody-based drug discovery and development of immunotherapies that harness the B-T cell axis.

## Introduction

Clear cell renal cell carcinoma (ccRCC) accounts for ∼80% of malignant renal tumours^1,2^. One third of patients present with metastatic disease at diagnosis, and recurrence occurs in 50-80% of cases after nephrectomy^1^. Alterations in the von Hippel Lindau (VHL) gene, hallmarks of ccRCC^3^, lead to dysregulated hypoxia-inducible factor (HIF) signalling and accumulation of HIF-2𝘢, which promotes angiogenesis^4,5^. Despite a relatively low mutational burden (1-2 mutations per megabase)^6,7^ ccRCC tumours are highly infiltrated by immune cells, particularly T cells^8^. The immune-rich, vascularised tumour microenvironment (TME) underpins the success of vascular endothelial growth factor receptor (VEGFR) tyrosine kinase inhibitors (TKIs) and, more recently, immune-checkpoint inhibitors (ICIs), both now standard-of-care treatments^9,10^. Small-molecule HIF-2𝘢 inhibitors have also shown promising clinical activity^11–13^. Yet, the 5-year survival rate for metastatic ccRCC remains at ∼10%^14^. A small subset of patients experience prolonged survival, often with cycles of disease stability and relapse^15,16^, suggesting ongoing immune-mediated tumour control^17^. Understanding the immune mechanisms underlying these exceptional outcomes could reveal biomarkers for improved risk stratification and therapeutic development. Tertiary lymphoid structures (TLSs) are immune cell clusters primarily composed of B and T cells, resembling germinal centres (GCs) of secondary lymphoid organs (SLOs)^18^. Their formation within the tumour microenvironment (TME) is influenced by the surrounding stroma^19^. Intratumoral TLSs are associated with improved survival and response to therapy across many cancer types^20^.

In ccRCC, most studies on anti-tumour responses have focused on CD8^+^ T cells. Clonal expansions, shared TCR clonotypes across patients^21^ and persistence of tumour specific T cell clones years after nephrectomy^22^ support a sustained immune response. However, the prognostic value of CD8^+^ T cells remains controversial^23,24^, and conventional risk models, developed before the use of ICIs, lack immune biomarkers^25^. B cells also play critical roles in cancer, either as antibody secreting cells (ASC), antigen presenting cells (APC) or through cell-cell signalling within the TME and lymphoid sites^26,27^. Memory B cells and plasma cells (PCs) are consistently linked to favorable outcomes^28,29^. In ccRCC, TLSs can support local B cell maturation and differentiation into PCs leading to tumour-bound antibodies and improved tumour control^30,31^. Recent studies highlight coordinated B and T cell responses within tumours. Intratumoral exhausted PD1^+^ CD8^+^ and PD-1^+^ CD4^+^ T cells express high levels of the CXCL13, a key B cell chemoattractant, favouring TLS formation^32,33^. Humoral responses in cancer may also generate autoantibodies against self-antigens, such as extracellular matrix (ECM) proteins^34^. Interestingly, autoreactive B cells, chronically stimulated, acquire somatic hypermutation (SHM) and shift BCR reactivity towards tumour antigens, a mechanism known as clonal redemption^35^.

Effective antitumour immunity depends on dynamic crosstalk between draining lymph nodes (dLNs) and TLSs. Pre-exhausted TCF-1^+^ PD-1^+^ CD8^+^ T cells originate in dLN GCs and migrate to the tumour, where they expand and further differentiate within TLSs. A key feature of the cancer immunity cycle^36^ is the dynamic interplay between tumour dLN and TLS, amplifying anti-tumour immune responses^37^. Similarly, clonal overlap of B cells between dLNs and tumour suggests active cycling across these sites^38^. Despite these insights, direct comparative studies of B and T cell clonal selection between dLN GCs and intratumoral TLS are limited^38^, and this interplay remains largely unexplored in ccRCC.

Here, we use a multimodal approach to characterise the 16-year longitudinal adaptive immune landscape of an exceptional long-term survivor with metastatic ccRCC. We demonstrate that persistent long-term immunity is not static but relies on coordinated clonal evolution and plasticity across sites. We identify a dense TLS network in the primary tumour, and coordinated clonal restriction of CD8^+^ T cells and B cells. Notably, we observe persistent B and CD8^+^ T cell clones throughout the disease course. Importantly, our findings highlight a role for TLSs in supporting clonal redemption of intrinsically autoreactive B cells through reiterated GC-TLS immune cycling, which fuels the generation and maturation of anti-tumour B cell responses. Finally, we identify tumour-reactive antibodies as potential candidates for antibody-based drug discovery.

## Results

### Clinical history of the metastatic ccRCC exceptional survivor

This study focuses on an exceptional long-term survivor of metastatic clear cell ccRCC, who demonstrated an overall survival of 16 years despite widespread metastatic disease. This provides a unique opportunity to study long-term tumour immune responses and drivers of exceptional tumour control.The patient was diagnosed with Stage III ccRCC (T3a, N0, M0) and underwent radical nephrectomy with local lymph node removal (year 0). Over the following years, multiple brain and bone metastases developed (**Supplementary Figure 1**), requiring surgery and radiotherapy (RT) and treatment with Zoledronic Acid and Sorafenib in years 2-5, achieving periods of stable disease. A thyroid metastasis was resected at year 7, followed by minor progression of bony metastases. At year 10, pancreatic metastasis emerged that stabilised after RT and later required total pancreatectomy, splenectomy, and lymph node removal at year 16. Importantly, the patient did not succumb to progressive ccRCC, but rather to complications arising post-operatively, underscoring the exceptional duration of successful tumour management.

### TLS features differ between primary and metastatic tumour sites

TLSs resemble germinal centres (GCs) of secondary lymphoid organs (SLO) and their density correlates with survival across cancers^28,31^. To explore how tumour location influences TLS formation and function, we compared immune infiltrates in primary ccRCC (year 0) and pancreatic metastasis (metPan, year 16), and phenotypic similarities between TLS and GCs in SLOs (ccRCC dLN year 0, metPan dLN and spleen, year 16) (**Figure 1a, Supplementary Table 1**). Using multiplex immunofluorescence (mIF), we quantified cell densities of CD20^+^ B cells, CD4^+^ and CD8^+^ T cells, CD68^+^ macrophages, and functional markers (HLA-DR, PD-1). TLS were defined as B cell follicles co-localised with T cells, and proximity to tumour cells was assessed via CA-IX staining.

**Figure 1.**
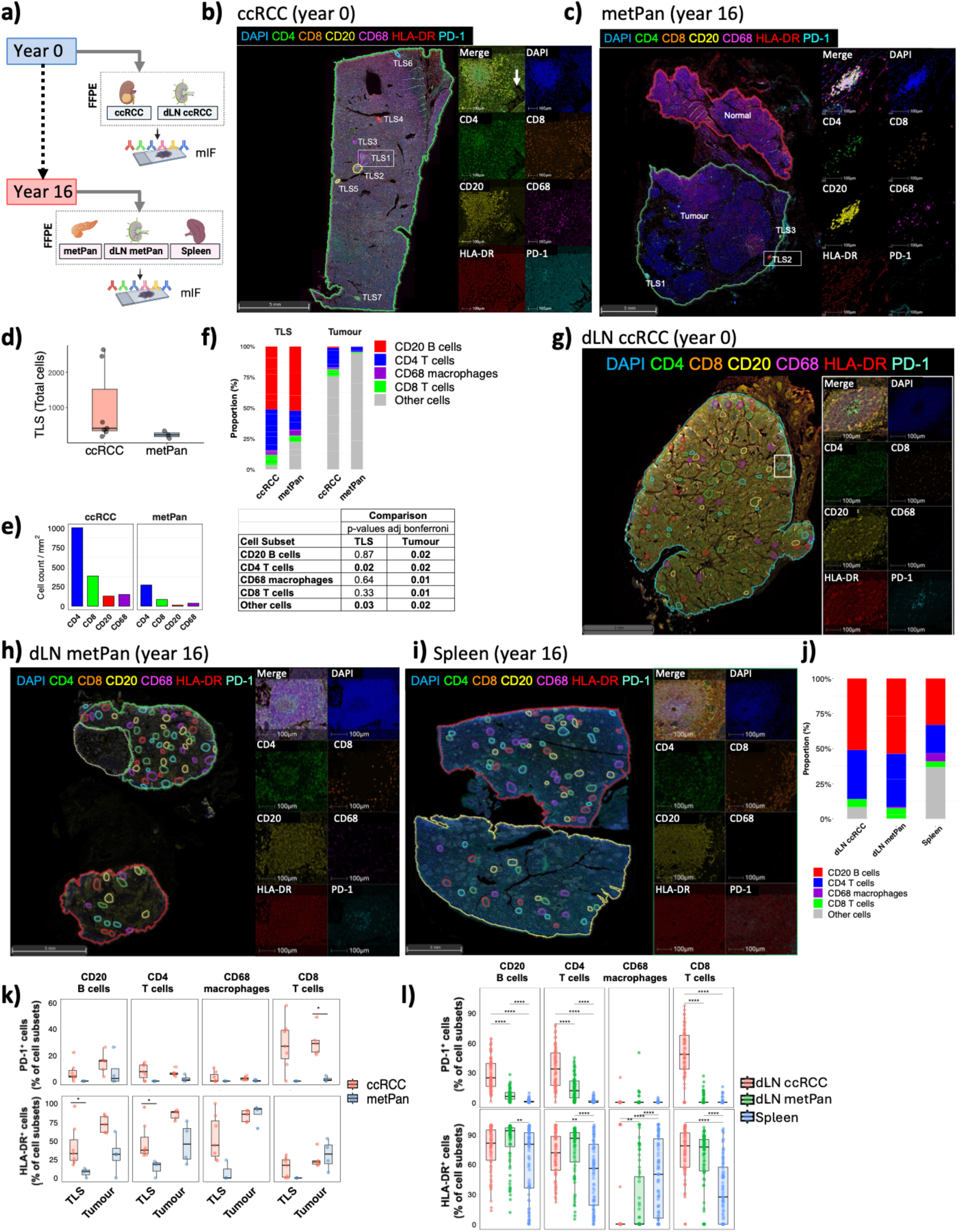
TLS differ between primary ccRCC and metPan and have comparable PD-1 expression profile with dLN GCs. a) Schematic representation of sample collection timepoints and experimental workflow. All images in Figure 1 were obtained by mIF on FFPE tissue sections using antibodies against CD4, CD8, CD20, CD68, PD-1 and HLA-DR, with DAPI for nuclear staining. b) Representative mIF image of the primary ccRCC tumour with annotated TLSs within the TME. The magnifications on the right show the expression of individual markers in the largest TLS (TLS1), egressing from a blood vessel (white arrow). c) Representative mIF image of the metPan with annotated tumour and adjacent normal areas, and TLSs in the peritumoral region. The magnifications on the right display the expression of individual markers in the largest TLS (TLS2). d) Boxplots showing TLS density and size (as total nuclei) within ccRCC and metPan (p= ns; permutation test, n = 10,000). e) Bar plots comparing immune cell content within tumour region of interest (ROIs) between ccRCC and metPan (cell count per mm^2^). f) Stacked bar plots showing the cellular composition of TLSs and tumour ROIs in ccRCC and metPan. (For each cell subset within TLS and tumour regions, the p-values of the comparison between ccRCC and metPan are shown in the table. Permutation test, n= 10,000). g) Representative mIF image of dLN ccRCC (year 0). For dLNs and the spleen, values refer to GCs, annotated in the images to represent immunologic structures comparable to TLSs. Magnified images on the right show individual markers in the highlighted GC. h) Representative mIF image of dLN metPan (year 16). i) Representative mIF image of the spleen (year 16). j) Stacked bar plots comparing the cellular composition across tumour dLNs (ccRCC and metPan) and the spleen, (Statistical Significance, **Supplementary Table 1**, Kruskal-Wallis test followed by Dunn’s test with Benjamini-Hochberg p-value adjustment). k) Boxplots showing the percentage of PD-1^+^ cells and HLA-DR^+^ cells (as % of cell subsets) within TLS and tumour ROIs in ccRCC and metPan (* p ≤ 0.05, Mann-Whitney test). l) Boxplots showing the percentage of PD-1^+^ cells and HLA-DR^+^ cells (as % of cell subsets) across GCs of dLNs and the spleen ( ** p ≤ 0.01, **** p ≤ 0.0001, Mann-Whitney test). For all TLS versus tumour comparisons, tumour areas were divided into multiple ROIs, excluding TLSs, as shown in Supplementary Figure 3a,b. (Abbreviations: TLS = tertiary lymphoid structure; CD4 = CD4 T cells, CD8 = CD8 T cells, CD20 = B cells, CD68 = macrophages; ccRCC = clear cell renal cell carcinoma; metPan = pancreatic metastasis; dLNs = tumour draining lymph nodes; GCs = germinal centres).

The primary ccRCC tumour contained numerous, large TLS distributed throughout the TME, while metPan displayed fewer, smaller TLS restricted to peritumoral areas. No TLS were found in adjacent normal pancreas (**Figure 1b-d; Supplementary Figure 3, Supplementary Table 2**). Sequential sections stained for CA-IX confirmed the close spatial association of TLS with tumour cells in ccRCC, in contrast to their complete exclusion from metPan CA-IX^+^ regions (**Supplementary Figure 2 a-b).** B cells comprised ∼55% of TLS immune cells at both sites, but ccRCC had significantly more CD4^+^ T cells than metPan (**Figure 1e-f**; **Supplementary Figure 4a**). SLO GCs showed comparable compositions, with higher macrophage density in the spleen (**Figure 1g-j**).

To assess the functional relatedness of immune niches, we compared the expression of PD-1, a key regulator of immune suppression^39^ and GC formation^40^ and HLA-DR, a marker of antigen presentation and cellular functionality^40^. PD-1^+^ cells were enriched in ccRCC TLS and dLN GCs, particularly among CD8^+^ T cells, but nearly absent in metPan and its dLN (**Figure 1k-l; Supplementary Figure 4c**). While PD-1 frequency varied across ccRCC TLSs, it did not correlate with TLS size (**Supplementary Figure 4b**). HLA-DR expression was lower in metPan TLSs than in ccRCC, suggesting reduced immune activation (Figure 1k-l; Supplementary Figure 4b). Interestingly, PD-1 and HLA-DR showed a negative correlation in ccRCC TLSs (p-value<1e-3), driven by an inverse relationship between PD-1^+^ CD8^+^ T cells and HLA-DR^+^ antigen-presenting cells (macrophages and B cells). This suggests that enhanced MHC class II antigen presentation may influence PD-1^+^ CD8^+^ T cell development in non-GC niches. In contrast, PD-1 and HLA-DR positively correlated in dLN GCs, with a slight opposite trend in the spleen (**Supplementary Figure 4d-e**).

These findings reveal site-specific differences in TLS formation. ccRCC harbored mature, functional TLSs throughout the TME, likely linked to active immune cycles with dLNs, whereas metPan exhibited immature peritumoral TLSs with limited activation. This further supports the functional coordination between tumour and dLNs, especially for CD8^+^ T cells^41,42^.

### Temporal persistence of B and T cell clones across sites

To understand the dynamics of long-term immune surveillance, we profiled B and T cell clonal architecture by combining single-cell and bulk BCR/TCR sequencing across sites and time points. This multi-platform approach allowed us to capture clonal information from both a recent blood and metastatic sample (via single-cell gene expression and BCR/TCR sequencing on FACS-purified CD45+ cells) and historical FFPE samples (via bulk BCR/TCR CDR3 sequencing) (Schematic in **Supplementary Figure 1b; Figure 2a**). The use of duplicate bulk sequencing runs enabled us to identify true clonal expansion with high confidence (**Supplementary Figures 5-6**).

**Figure 2.**
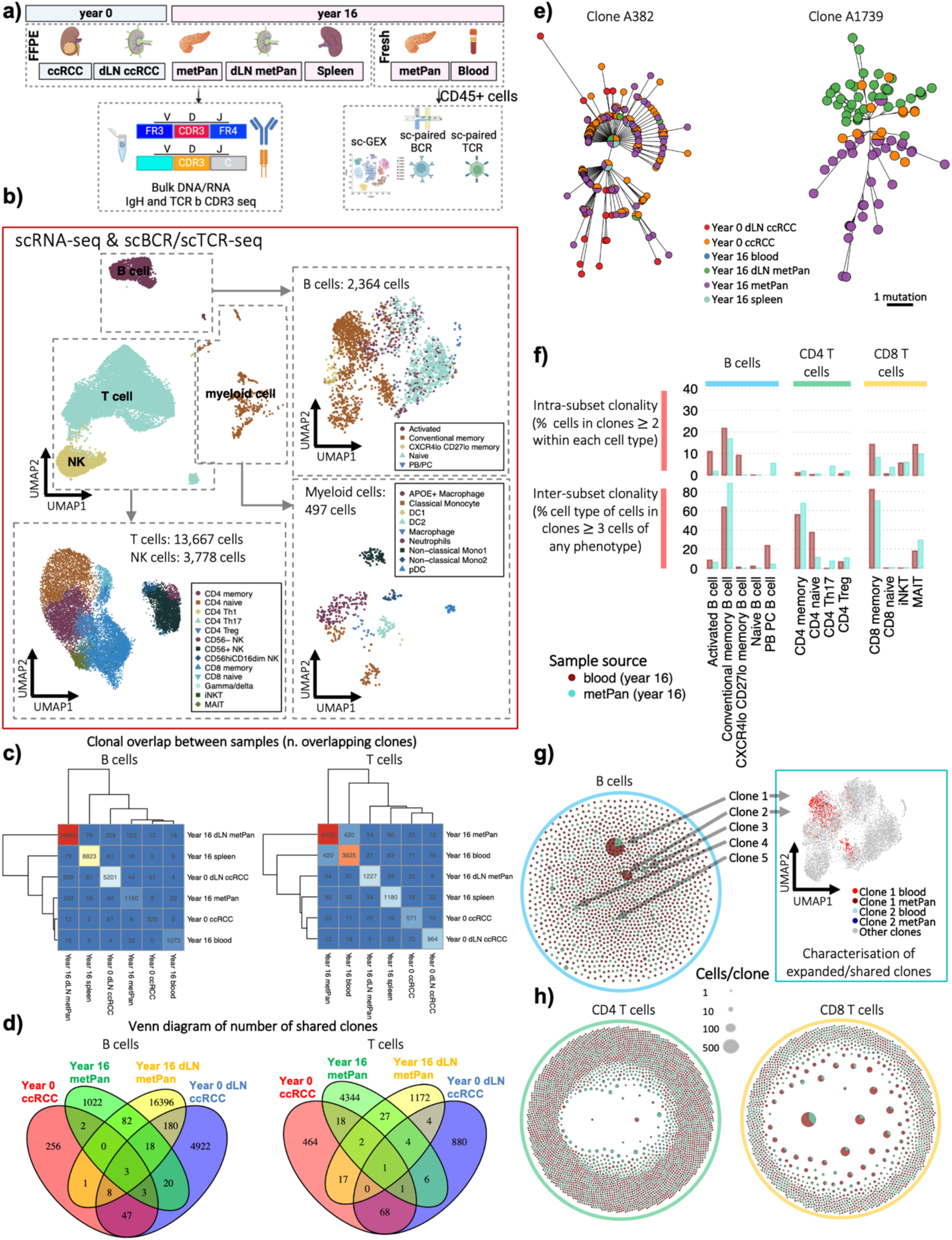
BCR and TCR clonal architecture across sites and time points. a) Schematic overview of the patient sampling strategy. RNA and DNA were available for year 0 ccRCC and its corresponding dLN and year 16 spleen, metPan and metPan dLN. CD45^+^ enriched cells from fresh metPan and blood were analysed by scRNA-seq with paired VDJ-seq. b) UMAP dimensionality reduction of the immune cells from the fresh CD45^+^ enriched metPan and blood samples depicting total immune cells (centre), B cells (right), myeloid cells (bottom right) and T and NK cells (bottom left). c) Clonal overlap (number of shared clones) between sites and time points for (left) BCRs and (right) TCRs. d) Venn diagram depicts B (left) and T (right) cells shared across sites at year 0 and year 16. e) BCR VDJ network plots showing two examples of persistent B cell clones detected at year 0 and year 16 and shared across multiple sites. These networks are based on maximum parsimony trees generated from BCR sequence alignments. f) Clonality of the B cell subpopulations from the scVDJ-seq between the blood and metPan via two measures: (top) *intra-subset clonality* (percentage of cells within clones containing >2 cells per subset, measuring the clonal expansion within specific subsets); and (bottom) *inter-subset clonality* (percentage of cells of each cell type within clones containing >3 cells across all populations, this indicates cells within each B cell subset that may be members of larger clones than span multiple phenotypes, reflecting B cell plasticity of expanding clones). g) VDJ clonality network plots for B cells. The insert shows the UMAP distribution for clonal members of clone 1 and clone 2. h) VDJ clonality network plots for CD4 T cells and CD8 T cells. In the clonality network plots, each node represents a unique VDJ sequence, and an edge represents a somatic hypermutation event (for B cells). Each node is represented by a pie chart showing the proportion of cells found in blood compared to metPan. The size of the nodes represents the number of cells expressing that VDJ sequence.

Single-cell analysis of CD45⁺ immune cells from year 16 metPan and blood identified the expected B, T, and myeloid populations^43^ (**Figure 2b**; **Supplementary Figure 5**). By integrating BCR and TCR sequencing data across longitudinal samples, we uncovered substantial B and T cell clonal sharing between tumour and dLNs, and remarkably, across the 16-year disease course and anatomical sites (**Figure 2c–d, Supplementary Figures 6b-c, 7**). This suggests ongoing clonal expansions and long-term maintenance across time and space. The highest levels of B cell and T cell clonal persistence were found between the year 0 ccRCC and its dLN with year 16 metPan (**Supplementary Figures 6b-c, 7b**). Furthermore, persistent B cell clones showed subclonal diversity, particularly within the dLNs, consistent with ongoing affinity maturation (**Figure 2e, Supplementary Figure 8a-b**). Four persistent B cell clones were captured by scRNA-seq year 16 metPan, and were found to be significantly enriched for memory and antibody secreting B cells compared to non-persistent B cell clones (Fisher’s exact test, p-value = 0.015, **Supplementary Figures 9a**). These findings are consistent with a continuous engagement of antigen-experienced memory B cells. Similarly, 166 and 118 T cells from persistent T cell clones were captured by scRNA-seq year 16 blood and Year 16 metPan, respectively, and were both found to be significantly enriched for CD8 memory T cells compared to non-persistent T cell clones (Fisher’s exact test, p-values <1e-6, **Supplementary Figure 9a**).

Altogether, these findings underscore the remarkable longevity and spatial coordination of both B and T cell responses in this patient, highlighting the sustained response of adaptive cells to chronic stimulation over extended periods of tumour evolution. The antigenic specificity and functions of these long-lived clones remains to be determined.

### Extensive B cell and CD8^+^ T cell clonal expansions between blood and metastatic pancreas

Single-cell RNA-seq revealed marked clonal expansions of both B and T cells in the blood and metastatic pancreas (metPan) at Year 16 (intra-subset clonality, **Figure 2f**). Over 10% of activated/memory B cells and CD8⁺ T cells belonged to expanded clones, with substantial clonal sharing between compartments (**Figure 2f-h, Supplementary Figure 9b**). T cell clones that were shared between the blood and metPan were significantly larger than site-restricted clones, consistent with a sustained systemic immune response (**Supplementary Figure 9c**). Expanded clones were predominantly CD8⁺ memory T cells (**Figure 2f**), enriched within the metPan compared to blood (**Supplementary Figure 9d**), and displayed preferential TRBV13 gene usage (**Supplementary Figure 9e**).

The two largest B cell clones (35.62% and 6.30% of blood B cells; 22.92% and 1.99% of metPan B cells), were IgD/M (unswitched) with conventional memory phenotype (**Figure 2g, Supplementary Figure 10a-c**). The largest clone (clone 1) used a mutated IGHV4-34 gene. IGHV4-34 autoreactivity is known to be driven by the FR1 AVY and CDR2 NHS motifs^44^. However, in clone 1, the FR1 motif mutated to AVF, suggesting clonal redemption through somatic hypermutation (**Supplementary Figure 10d**). These expanded B cell clones also upregulated antigen presentation signatures (HLA-DR and HLA-DPB1) compared with non-expanded intratumoural memory B cells (**Supplementary Figure 10e**). These findings align with prior observations in breast cancer, where B and T cell clonal evolution appears to co-track tumour neoantigen dynamics through MHC class II–mediated interactions^45^.Together, these data indicate a sustained, clonally focused adaptive immune response, shaped by chronic antigen exposure and tumour evolution.

### Intra-tumoural expanded B cells interact with CD8^+^ T cells in TLSs and GCs

Clonal expansion and trafficking of B and T cells between blood, tumours, and dLNs reflect anti-tumour activity^46–48^. To gain insight into the co-evolution of B and T cell clones in response to tumour antigens, we mapped two highly expanded B cell clones detected in both metPan and blood by scBCR seq (**Figure 3a**). To do this, we used *in situ* hybridisation on FFPE tissue with probes targeting the unique BCR heavy and light chain complementarity-determining region 3 (CDR3) of clone 1 (IGHV4-34) and clone 2 (IGHV3-72) (**Supplementary Figures 10b-c, 11; Supplementary Table 3**). A negative control dLN from an unrelated ccRCC patient was used to confirm probe specificity (**Supplementary Figure 11b**). Clone 1 B cells localised within a peritumoral TLS in year 16 metPan, and populated primary follicles and GCs of its dLN (**Figure 3b**). Clone 1 B cells were also detected in year 16 spleen and, to a lesser extent, in primary ccRCC TLS at year 0, indicating persistence over 16 years (**Supplementary Figure 12a, c, and d**). In contrast, clone 2 CDR3 transcripts were scarcely detected in year 16 metPan and its dLN, likely due to its smaller size (**Supplementary Figure 11c and d**). Multiplex immunofluorescence (IF) imaging on sequential year 16 metPan sections confirmed colocalisation of clone 1 B cells with CD4^+^ and CD8^+^ T cells within the peritumoural TLS (**Figure 3c**).

**Figure 3.**
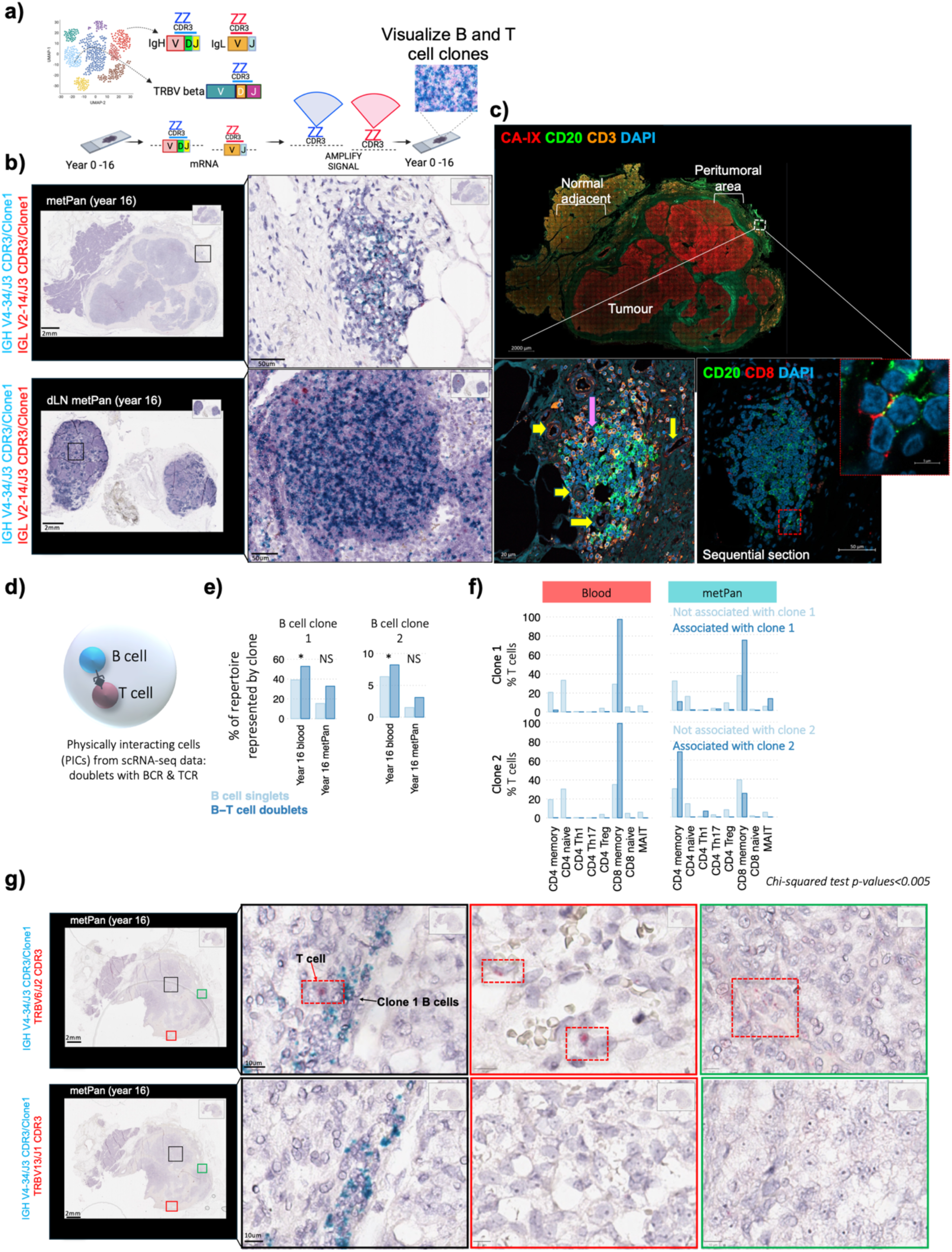
Tracking the spatial localisation of selected B and T clonotypes. a) Schematic of the Basescope assay. Probes specific to BCR and TCR CDR3 regions were designed from scBCR/TCR seq data to detect selected clonotypes in tissue. For B cells, probes targeted the heavy and light chain (IgH/IgL) CDR3 regions of Clone 1 (VH4-34) and Clone 2 (VH3-72); blue and red dots indicate IgH and IgL signals, respectively. For T cells, probes targeted the T cell receptor beta (TCR-β) CDR3 regions of Clone T1 (TRBV6) and Clone T2 (TRBV13). In the co-localization assay, the IgH probe for Clone 1 was combined with TCR-β probes for Clone T1 or T2, where blue dots indicate Clone 1 B cells and red dots indicate TRBV6 or TRBV13 T cells. b) Representative Basescope images of metPan (top) and dLN metPan (bottom) hybridised with Clone 1 B cells probes (IgH and IgL). Clone 1 V(D)J gene annotations are shown on the left. Clone 1 transcripts localised exclusively within a peritumoral TLS in metPan, and within GCs and follicles in the metPan dLN. Both heavy and light chains were detected, with stronger IgH signal (blue). Whole-section images, scale bar= 2 mm; magnified views, 50 µm. c) Representative mIF images of metPan using anti-CD20 and anti-CA-IX antibodies with anti-CD3 and, on sequential sections, with anti-CD8. These images correspond to the same TLS shown in the BaseScope assay. Staining highlights the co-localization of clone 1 B cells with CD3^+^ T cells and potential interactions with CD8^+^ T cells within the TLS. Pathologist annotations: yellow arrows indicate blood vessels, pink arrow indicates high endothelial venules (HEV). Whole section image scale bar= 2mm; magnifications, 20 µm (CD3) and 50 (CD8). d) Schematic showing a 10x Chromium droplet encapsulating a B cell and a T cell. Gene expression and VDJ sequencing of this droplet captures a hybrid transcriptome, revealing their physical interaction. e) Bar plot comparing the proportion of singlets (light blue) versus doublets (dark blue) for Clone 1 and Clone 2. Fisher’s exact test p-values are indicated with an asterisk (*) for statistical significance (p<0.05) or “NS” for not significant. f) Bar plots showing the proportion of T cell subsets that are clonally related to T cells found in doublets with Clone 1 B cells (top) or Clone 2 (bottom). This is compared to the proportion of T cells that are not doublet-related (light blue). g) Representative Basescope images performed on distal metPan sections, with distinct TLS density and distribution compared to those in panel ( c). V(D)J gene annotations are shown on the left. (Top) Clone 1 B cells (VH4-34) transcripts were detected within an intratumoral cluster with Clone T1 (TRB-V6) T cells located nearby (black rectangle), as well as within a peritumoral cluster and tumour nodes (red and green rectangles, respectively). Dashed red boxes highlight TRB-V6 T cells. (Bottom) No Clone T2 (TRBV13) signal was detected. Whole-section image, scale bar= 2 mm; magnifications, 10 µm. CA-IX = Carbonic Anhydrase 9, CD20 = B cells, CD3 = T cells, CD8 = cytotoxic T cells, and DAPI for nuclei. Basescope slides were acquired with Leica Aperio, 40X objective; mIF slides were acquired with the AiryScan confocal microscope, 20X objective, and magnifications with the 63X objective.

To assess direct B-T cell interactions, we analysed scRNA-seq sequencing data to identify droplets co-capturing B and T cells (doublets), defined by the presence of both a BCR and a TCR, with mixed B and T cell gene transcriptional profile (**Figure 3d**). Comparative analysis of doublet frequencies revealed that clone 1 and clone 2 B cells were significantly enriched in B-T doublets relative to their representation in the overall B cell population, suggesting preferential physical interactions with T cells compared to other B cell clones (**Figure 3e**). Phenotypic characterisation of T cells clonally related to the T cells found in doublets with clone 1 and clone 2 B cells demonstrated enrichment for CD8⁺ memory T cells compared to T cells not associated with doublets (p-values < 0.005, **Figure 3f**). Additionally, we observed preferential association with T cells within B-T cell doublets expressing TRBV13 gene segments (**Supplementary Figure 13**). These findings suggest that specific B and T cell clones exhibit non-random interaction patterns, potentially reflecting antigen-driven cognate pairing or shared tissue localisation preferences.

We next performed spatial mapping of clone 1 B cells in relation to two expanded T cell clones predicted to engage in cognate interactions based on doublet B-T cell co-capture: clone T1 (TRBV13) and clone T2 (TRBV6) on independent tissue sections. Clone 1 B cells were consistently identified within the Year 16 metPan tumour core (**Figure 3g**) as well as Year 0 ccRCC, dLN ccRCC (**Supplementary Figure 12a, and c**). Notably, clone T2 T cells exhibited a dispersed distribution across metPan regions, including direct spatial interaction with a clone 1 B cell (**Supplementary Figure 11**).

Together, these results demonstrate the long-term persistence of a highly expanded VH4-34 B cell clone across tumour, TLS, and secondary lymphoid sites for over 16 years, undergoing somatic hypermutation and maintaining spatial proximity to both CD4⁺ and CD8⁺ T cells. In contrast, expanded T cell clones displayed more transient and scattered residency, without clear enrichment in TLS or dLN GCs. The consistent detection of clone 1 B cells within tumour-associated structures and their preferential association with CD8⁺ T cells indicates selective and durable B-T cell interactions, supporting the concept of clonal redemption and suggesting a role in sustaining CD8^+^ T cell function.

### Tumour infiltrating B cell clones can specifically bind tumour cells

To assess whether expanded B cell clones recognised tumour antigens, we generated five monoclonal antibodies (Mab1-Mab5) from scBCR-seq. Mab1 and Mab2 corresponded to the two largest clones (Clone 1 and Clone 2), while Mab3-Mab5 were derived from metPan enriched clones (**Supplementary Figure 10a-c**). Germline revertants of Mab1 and Mab2 (Mab1-GL and Mab2-GL), lacking somatic hypermutation (SHM), were also produced to determine antigen-driven clonal evolution (**Figure 4a, Schematic**). Antibody reactivity was performed via IF on multiple epithelial cell lines, including RCC4 (clear cell renal cell carcinoma), HKC-8 (transformed kidney proximal tubule), and PANC-1 (pancreatic epithelioid carcinoma) and a non-tumour-derived hEM3 (human endometrial epithelial) (**Figure 4b and g; Supplementary Figure 14**). Anti-human N-Cadherin was used as a plasma membrane marker, with a isotype antibody and anti-CA-IX as negative and positive controls, respectively (Supplementary Figure 14c, d,e). Two staining strategies were used: fixed/permeabilised cells to detect intracellular epitopes, and live cells to identify extracellular or conformational epitopes.

**Figure 4.**
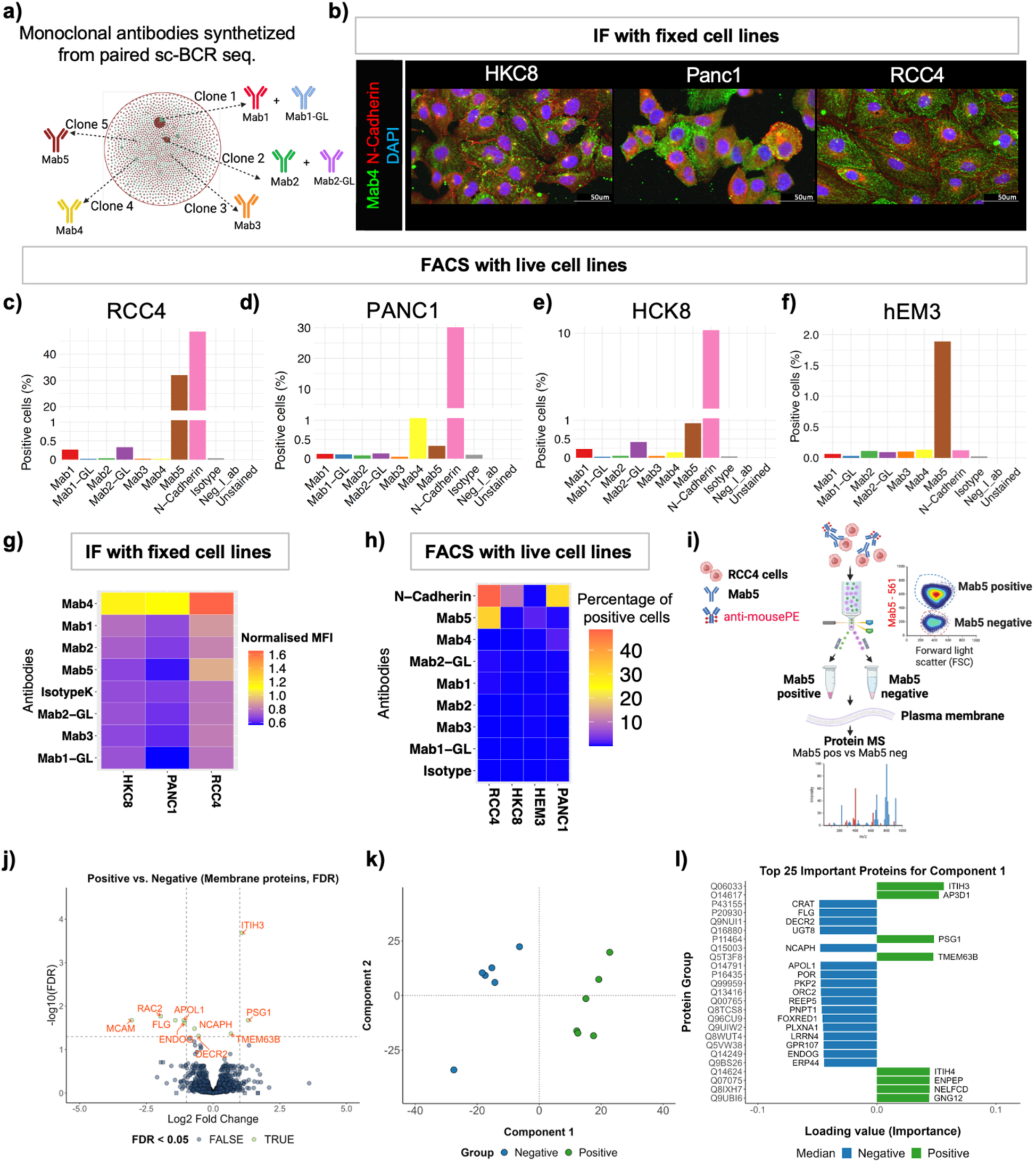
Monoclonal antibodies derived from circulating B cells exhibit high reactivity against RCC4 cells, with ITIH3 as a potential target. a) Schematic of monoclonal antibodies (Mab) synthetised from five selected B cell clones identified by scBCR-seq. Mab-1 and Mab-2 germline revertant (Mab1-GL and Mab2-GL) antibodies were generated from Mab-1 and Mab-2 by reverting all V(D)J positions to their germline counterparts. b) Representative IF images of three epithelial tumour cell lines (HKC8, PANC1, RCC4) stained with Mab4 (green) in combination with anti-N-Cadherin (membrane, red) and DAPI for nuclei. Cells were fixed and permeabilised with methanol. c) Flow cytometry analysis (FACS) analysis of live RCC4 cells. Bar plots show the percentage of positive cells for each Mab relative to N-Cadherin, isotype and negative controls. d) Flow cytometry analysis (FACS) analysis of live PANC1 cells. Bar plots show the percentage of positive cells for each Mab relative to N-Cadherin, isotype and negative controls. e) Flow cytometry analysis (FACS) analysis of live HCK8 cells. Bar plots show the percentage of positive cells for each Mab relative to N-Cadherin, isotype and negative controls. f) Flow cytometry analysis (FACS) analysis of live hEM3 cells. Bar plots show the percentage of positive cells for each Mab relative to N-Cadherin, isotype and negative controls. g) Heat map showing IF mean fluorescence intensity (MFI) for each Mabs across cell lines. Mab4 shows the highest reactivity, particularly against RCC4 cells. h) Heat map showing the percentage of FACS-positive cells across cell lines stained with monoclonal antibodies and anti-N-Cadherin. Mab5 displays significantly higher reactivity specifically towards RCC4 cells. i) Schematic of plasma membrane protein mass spectrometry (MS) used to identify potential Mab5 target antigens on RCC4 cells. j) Volcano plot of differentially expressed membrane proteins between Mab5-positive and -negative groups. Square points indicate proteins undetected in one group. Green colour represents proteins with FDR <0.05 (BH adjusted p value). k) Partial least square – discriminant analysis (DLS-DA) performed on log2 normalised protein intensities distinguishes Mab5-positive and -negative samples along Component 1. l) Top 25 proteins ranked by loading value stability score from sparse PLS-DA (sPLS-DA), indicating their contribution to sample separation along Component 1. Abbreviations: RCC4 = clear cell renal cell carcinoma; HKC8= human kidney proximal tubule carcinoma; PANC1 = pancreatic duct epithelioid carcinoma; hEM3 = human endometrial epithelial cells. Neg_I_ab= negative control, without the I unconjugated antibody.

Against fixed cells, Mab4 showed the strongest binding across tumour cell lines (p < 0.001) with preferential reactivity to RCC4 cells in a concentration-dependent manner via diffuse intracellular staining (**Figure 4b and g**; **Supplementary Figure 14a-b and c**), which was confirmed via an alternative permeabilisation method using PFA and Saponin (**Supplementary Figure 14f**). Mab5 showed weak binding to fixed RCC4 cells (**Figure 4g; Supplementary Figure 14a,b, d,e**).

In contrast, flow cytometry confirmed Mab5 as the most highly reactive BCR clone-derived antibody against live cells. It demonstrated robust binding to approximately 35% of RCC4 cells, a reactivity level comparable to the N-Cadherin control, while showing only minimal binding to irrelevant cell lines tested (hEM3, 1.89%; HKC-8, 0.19%; PANC-1, 0.33%) (**Figure4c-f, and h**). Mab5 also strongly bound dissociated autologous metPan tumour cells, with approximately 50% of CA-IX positive cells also staining positive for Mab5 (**Figure 5a-h**). Crucially, Mab5 exhibited consistent reactivity with unrelated ccRCC tumor cells but showed no binding to matched adjacent normal tissue (**Figure 5j; Supplementary Figure 15**), demonstrating tumour specificity.

**Figure 5.**
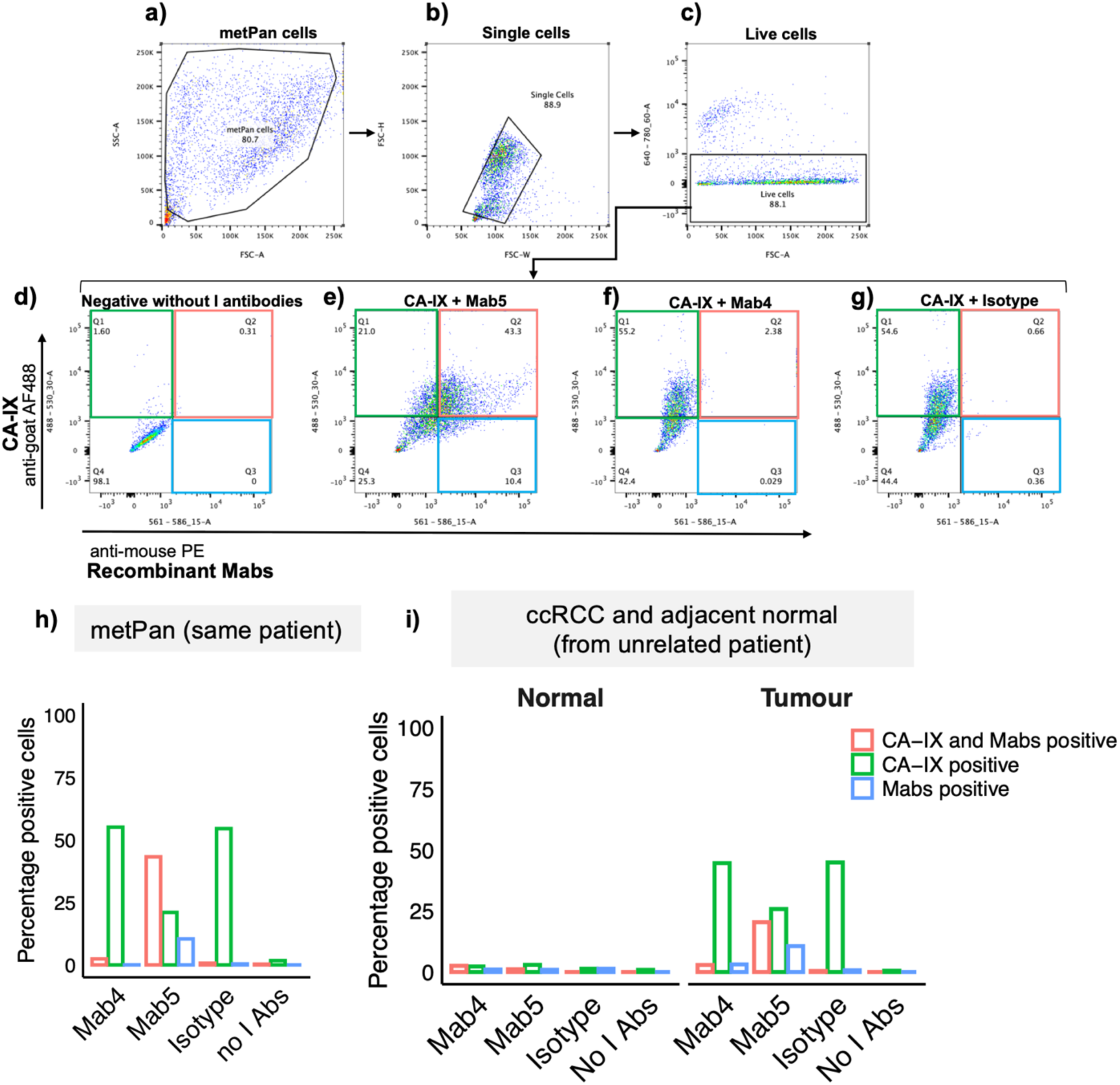
Flow cytometry analysis of monoclonal antibody reactivity against dissociated tumour cells from the patient’s pancreatic metastasis (metPan) and control tissue. The recombinant monoclonal antibody Mab5, derived from intratumoral B cells (clone 5), displayed tumour specificity, with strong binding to autologous metPan tumour cells and an unrelated ccRCC sample, but no reactivity toward adjacent normal tissue. Gating strategy for flow cytometry of dissociated metPan cells: a) tumour cells selected by forward scatter (FSC) and side scatter (SSC) parameters; b) doublets and aggregates excluded using FSC-W versus FSC-H; c) live cells identified using LIVE/DEAD™ Fixable Near-IR Dead Cell Stain; d) negative control without primary antibody included to establish background signal. e) metPan cells stained with anti-CA-IX (Carbonic Anhydrase 9) and Mab5; f) metPan cells stained with anti-CA-IX and Mab4; g) metPan cells stained with anti-CA-IX and mouse isotype control. h) quantification of recombinant monoclonal antibody binding to dissociated metPan cells. Bar plots show the percentage of antibody-bound cells for Mab4, Mab5, and isotype control, alongside anti-CA-IX antibodies. Due to limited cell count from metPan tissue, only Mab4 and Mab5 were tested. i) quantification of recombinant monoclonal antibody binding to dissociated cells from an unrelated ccRCC tumour and its normal adjacent tissue. Facet bar plots show the percentage of tumour and normal cells bound by each antibody (Mab4, Mab5, and isotype control), with anti-CA-IX antibody. Flow cytometric analysis was performed using a BD LSR Fortessa X-20 cytometer, and data was analysed with FlowJo software.

We additionally stained FFPE metPan tissue with the monoclonal antibodies, and the relative binding of tumour versus non-tumour regions was assessed. Mab1, Mab4 and Mab5 exhibited significantly higher binding to tumour cells compared to adjacent normal tissue in FFPE metPan tissue, but not in isotype controls (**Supplementary Figure 16a-c)**. Notably, when comparing Mab1 and its germline counterpart (Mab1-GL), only the somatically matured Mab1 preferentially bound the tumor regions within the FFPE metPan tissue (**Supplementary Figure 16d-f**). This finding suggests that SHM drove a shift in specificity from polyreactivity toward tumor-antigen recognition (**Supplementary Figure 16d-f**). Finally, to assess polyreactivity, all Mabs were tested for binding to healthy donor PBMCs. All antibodies showed minimal binding to these unrelated cells (<1.5%), with the notable exception of Mab1-GL (the unmutated germline sequence). Mab1-GL demonstrated a trend toward the highest binding, a finding consistent with the inherent autoreactivity associated with the germline IGHV4-34 gene segment (**Supplementary Figure 16g**).

Collectively, these results reveal diverse antigenic specificities among the circulating and tumour-infiltrating B cell clones, underscoring the functional complexity of the anti-tumour response. Specifically, Mab4 targets a diffuse intracellular tumour antigen, while Mab5 recognises a conformational “public” tumour antigen without detectable cross-reactivity to self-antigens. Most importantly, Mab1, derived from the intrinsically autoreactive IGHV4-34 B cell clone, gained tumour specificity through SHM, a phenomenon consistent with the mechanism of “clonal redemption.”

### ITIH3 as a potential target of B cell response in metastatic ccRCC

Target identification for Mab5 was achieved using a differential proteomics approach. RCC4 cells were stained with Mab5 and isolated into Mab5-positive and Mab5-negative fractions via FACS (**Figure 4i**; **Supplementary Figure 17a**). Plasma membrane proteins from each group (three replicates per group) were isolated and analysed by mass spectrometry (MS), with pooled samples included for comparison.

Across all samples, 3,330 membrane proteins of which 3,230 proteins were quantified in ≥ 50% of samples in at least one group (**Supplementary Figure 17b-c**). Differential expression analysis revealed ten significantly regulated proteins, three of which were enriched in Mab5-positive cells: Inter-Alpha-Trypsin Inhibitor Heavy Chain H3 (ITIH3), Pregnancy Specific Beta-1 glycoprotein (PSG1), and Transmembrane Protein 63B (TMEM63B) (FDR < 0.05, Wald test with Benjamini-Hochberg correction, **Figure 4j**). Partial least square-discriminant analysis (PLS-DA) clearly separated Mab5-positive and Mab5-negative profiles (**Figure 4k-l**, **Supplementary Figure 17d-f**), confirming ITIH3 as the top discriminant feature. Adaptor-related Protein Complex 3 Subunit Delta 1 (AP3D1), a protein involved in intracellular trafficking, also showed strong association and may influence the Mab5 reactive phenotype (**Supplementary Figure 17f**). Collectively these results identify ITIH3 as a candidate antigen for potent anti-tumour intra-tumoural B cell binding.

## Discussion

In this study, we comprehensively profiled the adaptive immune response of a metastatic ccRCC patient who survived 16 years post-diagnosis, using a multimodal strategy that integrated single-cell transcriptomics, antigen receptor sequencing, multiplex imaging, and antibody discovery. We identified a sustained and coordinated immune response characterised by high TLS density in the primary ccRCC tumour, extensive B and CD8^+^ T cell clonal expansion, ongoing clonal redemption of autoreactive B cells engaged in direct physical B-T cell interactions within TLSs, persistence of memory B and effector CD8^+^ T cells, and the presence of tumour-specific antibodies.

The spatial analysis of the TME revealed that TLS density and architecture were profoundly heterogeneous between the primary tumour and its metastasis. The primary ccRCC harboured numerous mature TLSs throughout the TME, whereas the pancreatic metastasis contained fewer, early-stage TLSs confined to peritumoral regions, indicating stromal constraints on TLS maturation. These observations align with studies linking TLS to improved survival in ccRCC^30,49^ and other cancers^38^. Despite expected genetic and transcriptional divergence of metastatic lesions^50^, our findings suggest that extrinsic stromal factors, rather than higher mutational burden, primarily drive TLS development^51,52^, as observed in other metastatic cancers^53^. Distinct PD-1 and HLA-DR expression patterns further distinguished the tumour sites. PD-1^+^ CD8^+^ T cells were enriched in primary ccRCC and its dLN, likely representing stem-like PD-1^+^ TCF-1^+^ precursor of cytotoxic effector cells that circulate between tumours and dLNs^54,55^ and are critical for tumour control and response to immune checkpoint blockade ^56^. The co-localisation of these cells with highly mature TLSs and elevated HLA-DR expression suggests that mature TLSs function as critical immunological hubs that continuously reinvigorate T cells recruited from the dLNs ^37,57^. Conversely, the scarcity of these PD-1+ precursors in the pancreatic metastasis, combined with the structural immaturity of its TLSs, suggests a disrupted T cell immune cycle and a compromised effector response at the metastatic pancreatic site. Collectively, the observed difference in immune architecture offers a profound insight into the patient’s therapeutic response pattern. The low density of mature, functional PD-1+ precursor cells in the metastatic TME suggests that checkpoint inhibition targeting the PD-1/PD-L1 axis would have limited efficacy due to a fundamental. The clinical observation that ccRCC patients with pancreatic metastases are often exquisitely sensitive to antiangiogenic agents but resistant to immune checkpoint inhibitors (ICIs) is supported by these spatial findings ^58^.

scBCR sequencing revealed expanded IgM^+^ memory B cells with SHM, including a large clone shared between blood and pancreatic metastasis that utilised the autoreactive IGHV4-34 gene^59^ SHM disrupted the autoreactive AVY motif, indicating clonal redemption. VH4-34 B cells were enriched in dLN and splenic GCs and were also detected in a peritumoral TLS of the metastasis, colocalising with CD4^+^ and CD8^+^ T cells. These redeemed B cells expressed high levels of HLA-DR and HLA-DP, supporting an antigen presenting function. In line with previous studies^60^, we observed close B-CD8^+^ T cell interactions within TLSs, and scRNA seq doublet analysis suggested direct physical associations between redeemed B cells and CD8^+^ T cells. Such B-CD8^+^ T cell crosstalk remains largely uncharacterised and may involve B cell MHC-I upregulation^61^, CD27/CD70 signalling^62^ or cytokine release^63^. These findings raise the possibility that BCR composition, autoreactive/redeemed versus non-autoreactive clones, may modulate CD8^+^ T cell fate, influencing the balance between tolerance and cytotoxicity^62–64^. The inverse correlation between PD1^+^ CD8^+^ T with HLA-DR^+^ APC in ccRCC TLSs suggests distinct APC-CD8^+^ T cell dynamics compared to SLO GCs.

Consistent with previous work^46,65–69^ we observed expanded CD8^+^ T cell clones shared between blood and metastasis, with enrichment for TRBV13. Spatial mapping of two expanded T cell clones (TRBV13 and TRBV6), both linked to IGHV4-34 B cells in doublet analysis, revealed heterogenous localisation throughout the pancreatic TME, with no enrichment in GCs or TLSs. These observations suggest distinct B and T cell clonal dynamics across lymphoid niches, with shorter T cell residency compared to B cells, and indicate that long-term re-educated B cells likely engage in heterotypic interactions with a diverse CD8^+^ T cell repertoire.

Our integrated single-cell and bulk antigen receptor sequencing revealed extensive B and T cell clonal sharing between tumours and dLNs, with reduced intratumoral diversity, consistent with findings in other cancers^70,71^. Persistent clones across disease progression included CD8^+^ effector T cells and IgM^+^ B cells with SHM. Together with evidence of clonal redemption, reduced class-switching and plasma cell differentiation, these findings suggest that chronic tumour antigen exposure sustains low-affinity, tolerant B cells that contribute to tumour control primarily via antigen presentation rather than antibody secretion. This aligns with recent work showing that blocking plasma cell differentiation while enhancing antigen presentation augments anti-tumour immunity^72^.

Finally, we identified tumour-specific antibodies derived from expanded B cell clones, providing a window of the antigenic landscape driving these coordinated responses. Two monoclonal antibodies, Mab4 and Mab5, derived from intratumoral clones, exhibited strong binding to RCC4 cells. Mab5, in particular, showed high specificity for conformational “public” tumour-antigens, without cross-reactivity to adjacent normal tissue. Proteomic profiling identified ITIH3 (Inter-alpha-trypsin inhibitor heavy chain 3) as a putative Mab5 target, highlighting a potential extra-cellular matrix-associated antigen driving humoral responses. Together, these findings provide insights into durable B and T cell responses in metastatic ccRCC and establish a framework for developing antibody-based or combination immunotherapies that harness the B-T cell axis.

## Materials and Methods

### Sample access

The patient underwent total nephrectomy for a stage III ccRCC in the left kidney. Tissue samples and the blood were collected under the Oxford Radcliffe biobank, project: 19/A098, reference: 19/SC/0173. Fresh ccRCC and normal adjacent tissue from an unrelated ccRCC patient were collected under reference 24/SC/0220. The unrelated patient had the following diagnosis: ccRCC, ISUP GRADE 4, pT3a pNX (UICC TNM 8th Ed.), excised. LEIBOVICH SCORE = 8 (HIGH RISK). Informed consent was obtained for all patients. The study was in strict compliance with all institutional ethical regulations. All samples were subjected to pathological re-review and histological confirmation by pathologists before analysis, and clinical information is provided in Results section.

### Preparation for scRNA-seq

Fresh sample collection, PBMC isolation and metPan digestion were performed as previously published^73^, and processed within 24 hours of resection. Briefly, pancreatic tissue was collected shortly after resection. Sample was initially mechanically disrupted using a scalpel into small pieces. The pieces were put into a 15 mL conical tube, with 9 mL of complete RMPI (10% FBS, 1% Pen/Strep and 1 mM Glutamine) and 1 mL of 10x hyaluronidase/collagenase solution (StemCell, 07912, Vancouver, BC, Canada). A first round of digestion was done at 37°C for 30 min in a pre-warmed shaker. The supernatant was collected without disrupting the tissue and a fresh digestion media was added (10 mL complete RPMI containing 200 U of collagenase IV (Lorne Laboratories, LS004194, Danhill, Berkshire, UK), 100 mL/mL of DNAaseI (Sigma, DN25, Gillingham, Dorset, UK) and 0.5 U of universal nuclease (Pierce, 88702, Waltham, MA, USA) for an additional 30 min of digestion as before. The supernatant was combined with the one from the first digestion step and the remaining tumour pieces were squeezed through a 100 mm tissue strainer with a further 10 mL of complete RPMI. The supernatants from all digestion steps were combined and centrifuged for 10 min at 300xg. Any residual red blood cells were removed with ACK solution. Cells were stained using a viability dye and anti-CD45 antibody diluted at 1:200 (Clone 2D1 PECy7, 368532, Biolegend) and sorted on an LSR Aria. PBMC were isolated from whole blood (9 mL) by density gradient centrifugation on Ficoll-Paque™ (Merck, GE17-1440-0). Cells were stained using a viability dye and anti-CD45 antibody diluted at 1:200 (Clone 2D1 PECy7, 368532, Biolegend) and sorted on an LSR Aria.

#### scMulti-omics sequencing and pre-processing

scRNAseq transcriptome (GEX), CITEseq (Cell Surface Protein), VDJ TCR and VDJ BCR profiling was performed using the Chromium 10x Genomics system. Chromium Single Cell Immune Profiling Reagent Kits v1.1 solution was used to capture and profile 500-10,000 individual cells per sample. Steps included; GEM generation, post GEM-generation clean-up, cDNA amplification, VDJ enrichment and library construction. Generated libraries included 10x Barcodes, which were used to associate individual reads back to the individual partitions. The library was sequenced using the Illumina NovaSeq 6000 platform. The analysis pipeline applied to process Chromium single-cell data to align reads and generate feature-barcode matrices was performed as previously described^74^. Briefly, gene expression FASTQ files were processed using Cellranger count (v3.1.0) to perform alignment, filtering, barcode counting, and UMI counting, using 10X Genomics’ GRCh38 v3.0.0 reference for Gene Expression analysis and IMGT’s reference for VDJ TCR and BCR analysis. It uses the Chromium cellular barcodes to generate feature-barcode matrices, determine clusters, and perform gene expression analysis.

#### Filtering, doublet detection and batch correction

For each sample, cells with fewer than 500 transcripts or 500 genes were filtered out. Normalisation and scaling was done using the standard Seurat pipeline. Principal component analysis (PCA) was performed on 5,000 highly variable genes (HVGs) to compute 50 principal components, then *Harmony* was performed (reference) for batch correction, UMAP for dimensionality reduction, and the Louvain algorithm was used for clustering. These clusters were then annotated broadly into B cell, T cell or myeloid clusters based on mapping of >10% BCR+ droplets and elevated CD19 expression, >10% TCR+ droplets and elevated CD3 expression, <10% BCR/TCR+ droplets, respectively.

Doublet identification and removal was performed using both DoubletFinder^75^ and MLtiplet^76^. Each cell type was subsetted into individual objects, and re-clustering within these objects was performed excluding genes which were likely to be influenced by experimental rather than biological factors^77^. These include genes encoding for TCR variable chain, ribosomal proteins, heat shock proteins, mitochondrial proteins, cell cycle proteins, HLA, and noise-related genes (MALAT1, JCHAIN, XIST). For the B and T cell objects, immunoglobulin variable, TCR variable and isotype genes were also excluded.

#### Cell type annotations

##### T/NK cell annotations

The re-dimensionality reduced T cell object resulted in 100 clusters generated by k-means. T cell clusters were defined as those with mean proportion TCR expression >0.3, with innate clusters being those with mean proportion TCR <0.3. Individual cells in innate clusters which expressed TRA or TRB sequences were labelled as NK-like T cells.

The innate cells were re-clustered without the T cells to generate 10 clusters, and were labelled by gene expression, ILC1 (*TBX21, IFNG, CCL3*), ILC3 (*RORC, AHR, IL23R IL1R1*), gdT (*TRDC*), NK (*EOMES*, *GZMA, GNLY, KLRC1*) based on *de Andrade, et al.*^78^. CD56 bright (immature) NK cells were labelled based on ADT-seq CD56 expression. The remaining NK clusters were labelled based on gene expression patterns to give phenotypic descriptions. NK transitional cells have greater expression of cytokines, chemokines and their receptors (*XCL1, XCL2, CXCR4*), NK mature cells have greater expression of cytotoxic genes (*GZMA, GZMB, PRF1*), NK terminal cells have greater expression of adaptive genes (*PRDM1*, *ZEB2*).

CD4 and CD8 clusters were defined by gene expression. As has been well documented in T cell single cell papers, there were clusters with overlapping CD4 and CD8 expression. Cells in overlapping clusters were reassigned at the single cell level if either CD4 or CD8 expression was higher. Memory phenotypes were label based on CD45RA, CD45RO, and CD62L expression. Naïve (CD45RA, CD62L), EMRA (CD45RA), EM (CD45RO), CM (CD45RO, CD62L). Further phenotypic labels were based on RNA expression. Exhausted (4 or more of the following: *HAVCR2, PDCD1, TOX, LAG3, CTLA4, TIGIT, CD38, ENTPD1*). CD4 cells: Treg (*FOXP3*), senescent (*B3GAT1, KLRG1, CD28-, CD27*-), Tfh (*BCL6, ICOS, CXCR5*), Th17 (*RORC*), Th2 (*GATA3*), Th1 (*TBX21*). Finally, clusters were labelled as activated based on HLA-DR ADT-seq expression.

##### B cell annotations

The re-dimensionality reduced B cell object resulted in 34 clusters generated by Louvain clustering, and AddModuleScore was used to identify enriched phenotypes. Plasma cells were defined as clusters with the percentage of droplets above the 95th percentile BCR nUMIs (percBCR_high) >40% and PC score>0.04, plasmablasts as percBCR_high >15%, naive B cells with >80% unmutated BCRs and >98% IGHD/M BCRs, and memory B cells with mean CD27 expression>0.1. The following cell types were based on AddModuleScores and mean gene expression: B cell memory activated (>0.3 activated score and *CD27* expression >0.1), B cell activated pre-memory (>0.4 activated score and *CD27* expression <0.1), B cell MZ (>0.8 FGR score and CD27 expression >0.1), B cell GZMB+ memory (*GZMB* expression>0.3 and *CD27* expression >0.1), B cell pre-GC (>0.2 GCB_FT or >0.02 preGC score), B cell GC (>0.3 GC score), of which B cell DZ GC (>0.9 DZ GC), B cell LZ GC (>0.3 LZ GC score). Finally, naive B cells were reassigned at the single cell level if there was >3 SHM, if the isotype was not *IGHD/M*, or if there was detectable *CD27* expression (activated memory) or without *CD27* expression (activated pre-memory).

##### Myeloid cell annotations

The re-dimensionality reduced myeloid subsetted object was used to identify enriched phenotypes. We down-sampled the cells to 2000 UMIs/cells and selected variable genes similarly to the seeding step of the clustering. To focus on biologically relevant gene-to-gene correlation, we calculated a Pearson correlation matrix between genes for each sample. For that purpose, expression values were log transformed (log(1+UMI(gene/cell))) while genes with less than 5 UMIs were excluded. Correlation matrices were averaged following z-transformation. The averaged z matrix was then transformed back to correlation coefficients. We grouped the genes into gene modules by complete linkage hierarchical clustering. Specifically, semi-supervised module analysis by complete linkage hierarchical clustering was carried out on variable, biologically-meaningful, and abundantly expressed genes^79^. For example, curated cell-cycle genes and other lateral programs (such as HLA-and HIST-) were excluded from module analysis. Subsequently, myeloid cells were assigned annotations at two levels of granularity based on prior knowledge of marker genes and modules, spanning PDAC and other cancer datasets.

#### BCR-seq/TCR-seq analysis

For the single cell data, we used the *scIsoTyper* pipeline to assign most probable BCR IGH and IGK/L chains per droplet (based on nUMIs) and most probable TCR TRA and TRB chains per droplet (based on nUMIs) from Sivakumar et al ^73^. For the single cell BCR data, a clone was defined as a combination of (a) group of B cells expressing BCRs that are connected by SHM (i.e. within a cluster in a BCR network, where edges are generated between BCRs differing by one mutation), and (b) group of B cells expressing BCRs with identical CDR3 region DNA sequences and identical IGHV and IGHJ family identities, as previously described^73^. This definition allows for capturing clonal B cells expressing BCRs that have undergone SHM both within the CDR3 region and outside. For the TCR data, cells sharing an identical TCR alpha and beta nucleotide CDR3 sequence were considered to be members of the same clone. *scClonetoire* ^73^ was used to quantify the intra- and inter-subset clonality and other repertoire metrics run on single cell multi-omics repertoire data. Intra-subset clonality measures the number of B, CD4 or CD8 T cell clones with 2 or more cells within each cell subset. This accounts for sampling depth differences between samples by generating a mean across 1,000 subsamples at a fixed depth for each sample (n=5 cells). Inter-subset clonality measures the percentage of B, CD4 or CD8 T cells of each cell type as members of clones of size 3 cells or more across all populations. This accounts for sampling depth differences between samples by generating a mean across 1,000 subsamples at a fixed depth for each sample (n=50 cells). These sampling depths were chosen to ensure values were captured across as many immune cell subsets as possible, even when the cell type was rare, whilst still ensuring representation across the sample.

#### Bulk BCR and TCR repertoire analysis

FFPE tissue blocks from each site (ccRCC, dLN ccRCC, metPan, dLN metPan, spleen) were cut into 10 µm thick sections. RNA was extracted with the High Pure FFPET RNA isolation kit (Roche, 06650775001) and DNA with the High Pure FFPET DNA isolation kit (Roche 06650767001), according to the manufacturer’s protocols. RNA and DNA were quantified using the Qubit RNA HS Assay kit (Invitrogen, Q32855) and the Qubit dsDNA HS Assay (Invitrogen, Q32854), respectively, with a Qubit 3.0 fluorometer (ThermoFisher Scientific).

Bulk B cell receptor (BCR) repertoire sequencing was performed on RNA from ccRCC, dLN ccRCC, metPan, and dLN metPan sections, using ∼200 ng total RNA per sample. First-strand cDNA synthesis was performed with SuperScript™IV Reverse Transcriptase (Invitrogen, 18090050) following the manufacturer’s instructions. A pool of five barcoded reverse primers targeting exon 1 of the constant regions of IgM, IgD, IgA, IgG, and IgE isotypes (final concentration 2 µM each) was used during reverse transcription. RT-PCR negative controls with no input RNA and positive controls with PBMC RNA were also included. The cDNA was treated with RNase H (Invitrogen, 18021-071), purified using AMPure XP beads at a ratio of 1.8x (Beckman Coulter, A63881), and eluted in 40 µL nuclease free H_2_O (Thermo Fisher scientific, AM9916).

All CDR3 amplifications were performed by two-step PCR using the 2x QIAGEN Multiplex PCR kit (QIAGEN, 206143). All PCR batches included positive and negative control tubes, with PBMC DNA and no template, respectively. For bulk RNA IgH CDR3 (Batch 2 in PCR schematic, Supplementary Figure 1b), the first-round PCR (100 µL total volume) used 20 µL purified cDNA with a mix of FR3-specific forward primers for IgH V genes (FR3, VH1-7) with 3’-JH reverse primers (3’-JH1, JH2, JH3, JH4, JH5, JH6). In one sample (RCC_LN_03_bc_FR3-CNUS), the reverse primer CNU (complementary to the RT-PCR reverse primer tag) was used. PCR cycling conditions were 95°C for 15 minutes, 30 cycles of 94°C for 30 sec., 63°C for 30 sec., 72°C for 1 min., and a final extension at 72°C for 10 min. Second-round PCR used 10 µL of the first-round product with FR3-specific forward primers and 5’-JH reverse primers (5’- JH1, JH2, JH3, JH4, JH5, JH6), under the same cycling conditions, but reduced to 25 cycles (IgH CDR3 primers adapted from Dongen J JM., et al. Leukemia 2003; Primer sequences in **Supplementary Tables 4-6**).

Bulk DNA IgH and TCRβ CDR3 PCRs were performed in duplicate (two replicates per sample) (Batch 3 in PCR schematic, **Supplementary Figure 1b**). The detection of the same CDR3 in both replicates was indicative of clonal expansion. DNA IgH CDR3 amplification followed the same two-step PCR protocol, using ∼ 400-1000 ng of DNA in the first-round PCR. For TCR β CDR3 amplification, ∼ 250-550 ng of DNA was used in the first-round PCR, with forward primers annealing to the TRB-V genes and reverse primers annealing to the 3’-UTR regions of each TRBJ primers. The second-round TCR β CDR3 PCR used 10 µL of the first-round product under the same cycling conditions, with TRB V specific forward primers and TRB J reverse primers. All primer sequences are provided in **Supplementary Tables 4-6**.

PCR products were size separated by agarose gel electrophoresis, followed by band excision and purification with the QIAquick Gel Extraction Kit (28704) according to the manufacturer’s instructions. Purified PCR products were quantified using the Qubit fluorometer.

For library preparation, purified CDR3 PCR products were indexed with the KAPA Hyper Prep kit (Roche KR0961 – v6.17) and KAPA Unique Dual-Indexed Adapter Kit (Roche, KK8726). Libraries were quantified using QUBIT dsDNA HS Assay (Invitrogen, Q32854), normalised to an equal molar concentration and pooled into a single library at >15 nM. Sequencing was performed on an Illumina NovaSeq X with a 2x300 bp configuration. Raw sequencing reads were filtered for base quality (median Phred score >32) using QUASR^80^. Forward and reverse reads were merged if they contained an identical overlapping region of >50bp, or otherwise discarded. Universal barcoded regions were identified in reads and orientated to read from V-primer to constant region primer. The barcoded region within each primer was identified and checked for conserved bases. Primers and constant regions were trimmed from each sequence, and sequences were retained only if there was >80% per base sequence similarity between all sequences obtained with the same barcode, otherwise discarded. The constant region allele with highest sequence similarity was identified by 10-mer matching to the reference constant region genes from the IMGT database^81^, and sequences were trimmed to give only the region of the sequence corresponding to the variable (VDJ) regions. Isotype usage information for each BCR was retained throughout the analysis hereafter. Sequences without complete reading frames and non-immunoglobulin sequences were removed and only reads with significant similarity to reference IGHV and J genes from the IMGT database using BLAST^82^ were retained. Ig gene usages and sequence annotation were performed in IMGT V-QUEST, where repertoire differences were performed by custom scripts in Python.

#### Clonal sharing analysis

Clonal overlap and sharing were assessed across heterogeneous immune receptor sequencing data, specifically combining single-cell VDJ sequencing (scVDJ-seq), bulk RNA VDJ sequencing, and bulk DNA VDJ sequencing datasets.

##### BCR Clonal Definition

BCR CDR3 regions were extracted and identified using the IMGT database. BCR clones were defined as groups of BCRs whose IGH CDR3 nucleotide sequences differed by a maximum of one nucleotide from any other clonal member and containing identical IGHV and J gene segments family usages (an extension of the MRDARCY method^83^).

##### TCR Clonal Definition

TCR clones were defined by an identical TCR beta chain CDR3 nucleotide sequence.

To account for the inherent differences in quantification across data types (scVDJ-seq UMI counts, bulk RNA VDJ-seq relative numbers, and bulk DNA VDJ-seq amplicon numbers), both BCR and TCR clone counts were normalised to a binary count matrix, with 1 and 0 representing presence or absence of clones, respectively. The number of unique clones shared between any two samples or time points was then computed along with the Jaccard overlap index

##### Clonal Diversity

The Gini index was calculated to measure clonal diversity. To ensure robust comparison, the index was determined from the mean of 100 iterative subsampling events, standardised to 442 unique BCR and 92 unique TCR CDR3 sequences per sample.

##### Doublet analysis

Droplets that contained two or more TCRs or BCRs that were clonotypically distinct were identified as doublets. Singlets were then screened to identify clones that were captured within the doublet populations. Proportions of cell phenotypes were calculated and Fisher exact test performed to determine if doublet-associated proportions were distinct from non-doublet-associated proportions.

#### Differential gene expression analysis and pathway analysis

Differential gene expression analysis was performed using Seurat FinaMarkers function. Differentially expressed genes were defined as adjusted p-values <0.05.

#### FFPE tissue specimens

Sequential 5-µm-thick formalin-fixed paraffin-embedded (FFPE) tissue sections were prepared from the following sites: primary ccRCC, dLN ccRCC, metPan, dLN metPan, and the spleen. FFPE sections were prepared for multiplex immunofluorescence (mIF) and *in situ* RNA hybridisation (BaseScope) assays. Additionally, tissue section scrolls were prepared for RNA and DNA isolation.

#### Multiplex immunofluorescence (mIF)

mIF was performed on 5-µm FFPE sections. Slides were baked at 60°C for 1 hour, and deparaffinised in xylene, rehydrated through ethanol (100%, 95%, and 70%), rinsed in ddH2O, and washed in 1X phosphate-buffered saline (PBS). Heat induced epitope retrieval (HIER) was performed in 1x sodium citrate solution (SCS; pH 6.0) using a TintoRetriever (Bio SB, Bioscience for the world) pressure cooker (low pressure for 11 minutes). Blocking was performed with 5% IgG-free bovine serum albumin (Ig-free BSA; Stratech) in PBS containing 0.2% Tween20 (PBS-T) for 1 hour at RT. Primary antibodies were diluted in 3% IgG-free BSA in PBS-T and incubated overnight at 4°C in a humidified chamber. After washing, secondary antibodies were incubated at 37°C for 45 minutes in a humidified chamber. The panel of antibodies used in each figure, including recombinant mouse monoclonal antibodies (Mabs), is detailed in **Supplementary Table 2**. Auto-fluorescence quenching was performed using Vector TRUEVIEW (Vector Laboratories, SP-8500) followed by mounting with Vectashield vibrance antifade mounting medium with DAPI (VWR, H-1800-2). Images were acquired on a AiryScan Zeiss confocal using ZEN v3.0 software.

Six-plex IF was performed on FFPE tissue using the OPAL™ protocol (AKOYA Biosciences, Marlborough, MA, USA) on the Leica BOND RXm autostainer (Leica, Microsystems, Milton Keynes, UK). Antibodies were: PD-1 (Cell Marque, Cat# 315M-95) - Opal 480, CD4 (Leica, Cat# NCL-L-CD4-368) - Opal 520, CD20 (Dako, Cat# M0755) - Opal 570, CD8 (Dako, Cat# M7103) - Opal 620, HLA-DR (Santa Cruz, Cat# sc53319)- Opal 690, CD68 (Dako, Cat# M0876) - Opal 780, and DAPI for nuclei staining. Primary antibodies were incubated for 1 hour and detected with the BOND™ Polymer Refine Detection System (DS9800, Leica Biosystems, Milton Keynes, UK) as per manufacturer’s instructions, substituting DAB with Opal fluorophores. Antigen retrieval was performed prior to each antibody application using Epitope Retrieval (ER) Solution 2 (AR9640, Leica Biosystems) at 100°C for 20 minutes, following the standard Leica protocol. Slides were mounted with VECTASHIELD^®^ Vibrance™ Antifade Mounting Medium with DAPI (H-1800-10, Vector Laboratories, Burlingame, CA, USA). Whole slide and multispectral images (MSI) were acquired on the AKOYA PhenoImager and analysed with HALO software (Indica Labs).

#### Epithelial cell lines

RCC4 (clear cell renal cell carcinoma), HKC-8 (transformed kidney proximal tubule), PANC-1 (pancreatic epithelioid carcinoma), and a non-tumour-derived hEM3 (human endometrial epithelial) were grown in complete medium with Dulbecco’s Modified Eagle’s Medium High-Glucose (Sigma, D5796) supplemented with 10% foetal calf serum (FCS) (Sigma, F2442) and 1% penicillin/streptomycin (Gibco, 15140-122). Cell lines were grown in 175 cm^2^ flasks (Nunc, 159910) in an incubator at 37°C and 5% CO_2_.

#### Immunofluorescence (IF) using mouse monoclonal antibodies (Mabs) and cell lines

Cells were cultured in 175 cm^2^ flasks until they reached 80-90% confluency, washed with D-PBS 1X (Merck, D8537), and detached using TrypLE (Gibco, 12604013). After neutralization, cells were pelleted and washed in D-PBS 1X , and resuspended at a final concentration of 50 cells/µL. Approximately 10^4^ cells were seeded per well in a µ-Plate 96 well Black ibiTreat plate (ibidi 89626), and incubated overnight at 37°C in a CO_2_ incubator. Cells were fixed with cold methanol, washed with PBS 1X, and with 5% IgG-free BSA and 0.2% Tween20 in PBS 1X (PBS-T). Cells were then incubated with recombinant mouse monoclonal antibodies (Mabs) at varying concentrations (5 µg/mL, 10 µg/mL, 20 µg/mL) and anti-h N-Cadherin at 10 µg/mL (anti-h CD325, R&D System AF6426). Anti-human CA-IX (abcam, ab15086) and a mouse isotype control (Mouse Isotype kappa Invitrogen 14-4714-82) were used as controls. Primary antibody incubation was overnight at +4°C. Cells were then washed and incubated with secondary antibodies: anti-mouse Alexa Fluor 488 (Invitrogen #A32766) and anti-sheep Alexa Fluor 594 (Invitrogen A-11016). For the positive control well, anti-sheep Alexa Fluor 594 and anti-rabbit Alexa Fluor 488 (Invitrogen A32790) were used. Secondary antibodies were incubated at +37°C for 45 minutes. Cells were counterstained with Ibidi Mounting Medium with DAPI (Thistle Scientific, IB-50011). Images were acquired using the CSU-W1 SoRa (Olympus) spinning disk confocal microscope (Olympus) and analysed using Arivis software. For comparison, IF was also performed on PFA-fixed cells (4% paraformaldehyde) (Fischer Scientific 28908) and 0.1% Saponin (Merck, SAE0073).

#### Flow Cytometry analysis using cell lines and recombinant mouse monoclonal antibodies

Cells were cultured in 175 cm^2^ flasks to 80-90% confluency, washed with D-PBS 1X and detached using TrypLE. After neutralization with complete medium, cells were pelletted, washed, and resuspended in PBS 1X at a final concentration of ∼5 x10^5^ /mL. Cells were transferred into FACS tubes. Dead cell discrimination was performed using LIVE/DEAD™ Fixable Far Red Dead Cell Stain Kit (ThermoFisher Scientific, L34974) following the manufacturer’s instructions. After staining, cells were washed with PBS 1X and resuspended in blocking buffer (2% BSA in PBS 1X; Bovine serum albumin, Fisher Scientific BP1605-100). Cells were then incubated with mouse monoclonal antibodies (Mabs) at a final concentration of 10 µg/mL each. Positive control tubes were prepared using anti-human N-Cadherin ( anti-h CD325, R&D System AF6426), while negative controls included samples with no primary antibodies and samples stained with mouse IgG1 isotype control (clone X40, BD Biosciences, 349040). Primary antibody incubation was performed for 20 minutes on ice. Following primary antibody staining, cells were washed in 2% BSA in PBS 1X and incubated with secondary antibodies, Rat anti-mouse PE IgG1 (BD Pharmingen Clone A85-1/#550083) and the positive control tube with Donkey anti-Sheep Alexa Fluor 594 (Invitrogen A-11016). Cells were incubated for 20 minutes with secondary antibodies on ice and protected from light. Cells were then washed with PBS 1X and resuspended in 2%BSA in PBS1X. Samples were acquired using a BD Fortessa X20 flow cytometer, and data was analysed using FlowJo software.

#### Flow cytometry analysis of dissociated pancreatic metastasis (metPan) and recombinant mouse monoclonal antibodies

Fresh pancreatic metastasis (metPan) from the exceptional LTS was dissociated as described above, and the resulting single cell suspension was cryopreserved in FBS 10% DMSO (Thermo Fisher Scientific, J66650.AK). Fresh ccRCC and adjacent normal tissue samples, from an unrelated ccRCC patient, were collected soon after surgical removal, placed into CUSTODIOL® (HTK Solution) (PHARMAPAL) and kept refrigerated to prevent tissue degradation during transportation to the lab. Samples were transferred into Petri dishes, washed with DMEM to remove residual Custodiol solution, and then transferred into new Petri dishes containing 1mL (or less) of digestion medium. Tissues were minced using sterile scalpels and transferred into 10 mL of digestion medium. Digestion medium consisted of DMEM + Collagenase/Dispase (cat no. 10269638001, Merck Life Science, UK Limited) 1 mg/mL final concentration, Dnase I (Roche, cat no.10104159001) to 5 µg/mL final concentration. Tubes were incubated at 37°C for 30 minutes in pre-warmed orbital shaker. The supernatant was collected by filtering through a 100 µm cell strainer mounted on a 50 mL falcon tube. To stop enzymatic activity, 10 ml of complete medium (DMEM + 10% FBS) was added to the filtered suspension (through the strainer to further collect cells), which was then kept on ice.

For the remaining undigested tissue, an additional 10 mL of fresh digestion medium was added, and tubes were incubated for a further 30 minutes at 37°C with shaking. After the second digestion, the resulting cell suspension was filtered through the same 100 µm strainer and added to the previous supernatants. Cell suspensions were washed by adding complete medium and centrifuged at 300xg for 5 minutes. Cell pellets were washed and resuspended in cryopreserving medium (FBS 10% DMSO). For flow cytometry, cell suspensions were thawed in a 37°C water bath and immediately resuspended in pre-warmed Advanced DMEM/F12 medium (Gibco, 12634010) supplemented with 1% Glutamax (Gibco, 35050061). Cells were centrifuged at 300xg for 5 minutes at +15 °C, washed with PBS 1X, and centrifuged at 300xg for 5 minutes at +15°C, and the cell pellet was resuspended in 1 mL PBS 1X. Cell count and viability were assessed immediately using 0.4% Trypan blue (Gibco, 15250061) and a haemocytometer. Dead cell discrimination was performed using LIVE/DEAD™ Fixable Near-IR Dead Cell Stain Kit (Life technologies, L10119A) following manufacturer instructions. Cells were incubated on ice for 20 minutes, washed in 2% BSA in PBS1X and resuspended in 200 µL of the same buffer per FACS tube.

Cells were stained with anti-CA-IX antibody (R&D Bio-Techne AF2188) at a final concentration of 0.25 µg/tube and one of the Mabs (Mab1, Mab4 and Mab5) at a final concentration of 20 µg/mL. Negative control tubes included cells with no primary antibodies and cells stained with a mouse IgG1 Isotype control (clone X40, BD Biosciences, 349040) at the same concentrations as Mabs. Cells were incubated with primary antibodies for 20 minutes on ice. After incubation, cells were washed with 2% BSA in PBS1X and stained with secondary antibodies: rat anti-mouse PE (Clone A85-1, BDPharmingen, 550083) 5 µL/tube and Donkey anti-goat (1:400 dilution) (Invitrogen, A11055). Cells were then washed and resuspended in 200 µL of 2% BSA in PBS1X for acquisition. Samples were acquired on a BD Fortessa X20 flow cytometer, and data was analysed using FlowJo software.

#### Cell sorting of Mab5-positive RCC4 cells and plasma membrane protein isolation

RCC4 cells were cultured in 175 cm^2^ flasks and detached using TryplE (Gibco, 12604013) when they reached 80-90% confluency. After detachment, cells were washed and resuspended in PBS 1X. An aliquot was used to count cells using 0.4% Trypan Blue. Cells were then aliquoted into FACS tubes at a density of ∼6 x 10^6^ cells per tube.

Cell viability was assessed using Live/Dead Far Red (ThermoFisher, L34974), following the manufacturer’s instructions. After washing, cells were stained with the recombinant mouse monoclonal antibody Mab5 (final concentration 20 µg/mL), washed, and incubated the secondary anti-mouse PE (5µL/tube) (Clone A85-1, BD Pharmingen, 550083). After staining and washing, cells were resuspended in 2% BSA in PBS 1X and acquired on a BD FACSAria^TM^ III cell sorter. Mab5-positive and Mab5-negative populations were established based on gating established using a negative control sample (omitting the primary antibody, Mab5). These two cell groups were sorted into separate tubes for downstream processing. Sorted cells were centrifuged at 300xg for 5 minutes and the pellets were processed for plasma membrane protein isolation using the Mem-PER Plus Membrane Protein Extraction Kit (Pierce, 89842). Halt™ Protease and Phosphatase Inhibitor Cocktail (100X, Thermo Scientific, 78440) was added to both permeabilization and solubilisation buffers. Briefly, sorted cells were washed in 3 mL of the supplied wash buffer and centrifuged at 300x g for 5 minutes. Pellets were then resuspended in 0.75 mL of permeabilization buffer, vortexed briefly and incubated at 4°C for 10 minutes with constant mixing. Following centrifugation at 16,000x g at 4°C for 15 minutes, the supernatants containing cytosolic proteins were transferred into clean tubes. The pellets were resuspended in 0.5 mL of solubilisation buffer and incubated at 4°C for 30 min with constant mixing, and centrifuged at 16,000x g for 15 min at 4°C. The supernatant containing solubilised membrane proteins was transferred to clean tubes, and protein concentration was determined using the BCA Protein Assay kit (Pierce, 23227). We determined that ∼1 x 10^6^ sorted cells yielded approximately 20µg of plasma membrane proteins. A total of 12 samples were prepared, comprising six replicates each for Mab5-positive and Mab-5 negative populations. For each sample, both cytosolic and plasma membrane protein fractions were isolated, with a slight modification: following the permeabilization step, pellets were resuspended in 120 µL of 1% SDS and vortexed prior to submission. All samples were submitted to Inoviv for mass spectrometry (MS) analysis.

#### Mass spectrometry (MS)

After checking the total protein concentration with Pierce Micro BSA kit (quantified in triplicate), 10 μl from each of the 12 samples (Mab5-positive + Mab5-negative) was taken to form a general pool. Nineteen μl of each 12 samples and 4 aliquots of the general pooled samples were taken and mixed with 4 μl of 20% SDS Lysis buffer for S-Trap processing where 5 μg of trypsin was added to each sample and digestion was allowed to occur at 47 °C for 90 minutes^84^. Tryptic peptides were eluted from the S-Trap then dried down.

##### Label-free Discovery Proteomics

Samples were re-suspended in 3% ACN/0.5 % FA ready for analysis. Nano-LC-MS/MS label-free Discovery Proteomics analysis of the Mab5 positive and Mab5-negative RCC4 plasma membrane samples plus pooled controls was carried out using low pH reverse phase chromatography on a Thermo Ultimate 3000 Rapid Separation LC (RSLC) nano LC system coupled to a Thermo Exploris 480 Orbitrap. Samples were first loaded onto a trapping column: Acclaim PepMap100 C18 LC column (5 mm Å∼ 0.3 mm i.d., 5 μm, 100 Å, Thermo Fisher Scientific) at a flow rate of 10 μL min−1 maintained at 45°C followed by analytical separation on a 200 cm μPAC C18 nanoflow column with a bed width of 315 μm and a pillar height of 18 μm at a flow rate of 400 nl/min over a 120 min linear gradient (175 min overall run time). (3%-35%B; B is 80% Acetonitrile, 0.05% Formic acid). One µg of protein digest from each sample was loaded on column for each Nano-LC-MS/MS run. Samples were analysed on a Thermo Exploris 480 Orbitrap operated in Data Independent Acquisition mode. For DIA acquisition in full scan mode, the MS1 resolution was set to 45,000, with a normalised automatic gain control (AGC) of 300%, dynamic maximum injection time. DIA MS/MS spectra were acquired with 40 variable width windows covering 390 - 1621 m/z, with MS2 resolution set to 20,000, dynamic maximum injection time with an AGC target of 300%, and NCE of 30%.

##### Data Processing

The raw mass spectrometry data has been processed with: DIA-NN version 1.8^85^. Data was searched against human proteome database (Uniprot UP000005640) combined with common Repository of Adventitious Proteins (cRAP). Settings Summary: Fragment m/z: 200-1800; Enzyme: Trypsin; allowed missed-cleavages: 1; peptide length: 7-30; precursor m/z 300-1800; precursor charge: 1-4; Fixed modifications: Carbamidomethylation (Cys); Variable modifications: Oxidation (Met), Acetylation (N-term).

##### Data Analysis

Data were imported into proLFQua^86^ for normalisation and statistical analysis. Peptide data were summarised to protein-level by robust summarization and normalised by robust scaling. Proteins included those with <50% coverage in either sample group to allow proteins tahr were completely missing from one sample group to be analysed. Proteins that were completely missing in either group were imputed using random draws from a Gaussian distribution around the 0.1th percentile within the samples they were missing and used to estimate the sample group average in the comparison.

Antibody-derived proteins (i.e., heavy and light chain immunoglobulins) were excluded from the analysis. Membrane and cytosolic proteins were analysed separately. Principal component analysis (PCA), Spearman correlation analysis and hierarchical clustering were carried out to visualise sample group differentiation. Statistically significant differentially expressed proteins were identified through linear modelling in the proLFQua framework. Fold changes were calculated and plotted against false discovery rate (FDR) and raw p.values. Thresholds used for significant differential expression were defined as FDR ≤ 0.05. In absence of significance using FDR, significance was assessed using raw p.value < 0.05 with the caveat that this may contain a high number of false positives. Partial Least Squares Discriminant Analysis (PLS-DA) and sparse PLS-DA models were created using mixOmics^87^.

Functional enrichment analysis was carried out on selected proteins using Clusterprofiler^88^ against Gene Ontology, KEGG Pathways and Reactome Pathways. Membrane protein enrichment analysis was performed using the AFTM^89^ and MatrisomeDB^90^ databases. A total of 3330 membrane proteins were identified and 2380 cytosolic proteins were quantified with 2584 membrane proteins (or membrane-associated proteins) and 1690 cytosolic proteins present across all samples (**Supplementary Figure 17c**). 3230 membrane proteins and 2264 cytosolic proteins that have at least 50% coverage in either positive or negative groups were taken forward for further analysis.

#### BaseScope assay on FFPE tissue with probes directed against IgH and TCRβ CDR3 regions

The BasesScope assay was performed using the BaseScope^TM^ Duplex Detection probes (ACD) kit, according to the manufacturer’s manual. The assay consisted of target probes binding mRNAs of interest and a signal amplification system (Schematic in **Figure 3a**). The first BaseScope assay was performed to detect the two largest B cell clones shared between the peripheral blood (PB) and the pancreatic metastasis (metPan) identified by scBCR-seq, clone 1 (IgH V4-34/D2-21/J3; IgL V2-14/J2) and clone 2 (IgH V3-72/D1-26/J4; IgK V3-20/J2). The probes were specific for the IgH and IgL CDR3 regions, and were used on FFPE tissue from ccRCC, dLN ccRCC, metPan, dLN metPan.

The detection of green and red dots in the same cell, indicated that Clone 1 (IGHV4-34) and/or Clone 2 B cells (IGHV3-72) were present in the tissue. Positive control probes targeting housekeeping (HK) genes PPIB and POLR2A were used to assess integrity of the mRNA molecules (staining performed on sequential slides). To test their specificity, Clone 1 and Clone 2 BCR probes were also used on tumour LN tissue obtained from a non-related patient with metastatic ccRCC. After staining with the probes, slides were counterstained with 50% haematoxylin and mounted with VectaMount (Vector Laboratories) mounting medium. Images were acquired using the Aperio ImageScope pathology slide viewing software (Leica) with 40x objective. The second BaseScope assay was performed following the same protocol and was designed to detect Clone 1 IgH CDR3 (IGHV4-34) and the TCRβ CDR3 of two T cell clones (clone T1 TRBV13 [TRBV13*01 TRBD1*01 TRBJ1*01] and clone T2 TRBV6 [TRBV6-1*01 TRBD1*01 TRBJ2-5*01]) identified with sc-TCR-seq. This assay was performed on all FFPE sites, including the spleen. The schematic of probe combination per tissue sample is shown in **Supplementary Table 3**.

## Code and data availability

All code is available via https://github.com/rbr1/RCC_metPan. Data will be made available via _XX_ (currently in progress).

## Acknowledgements

Firstly, we would like to thank the patients and clinicians who contributed to this study. R.B.-R. and F.A.T. were supported by the Wellcome Trust, University of Oxford and Oxford Cancer Centre. S.A. was funded by Clarendon in partnership with St John’s College. We thank Theodosios Kyriakou and Angela Lee for overseeing the sample submission at the Centre for Human Genetics in Oxford. The research was supported by the Wellcome Trust Core Award Grant Number 203141/Z/16/Z with additional support from the EPA Cephalosporin Fund (CF 390). We thank the Inoviv Team for RCC4 plasma membrane protein processing and MS analysis. We thank David Church and Luciana Gneo for hEM3 (human endometrial epithelial) epithelial cell lines, Eric O’Neill for the PANC1 (Pancreatic cell line) and Daniela Moralli for insightful guidance on IF staining.

## Authors’ contributions

S.S., F.T., and R.B-R conceived and designed the analysis. E.D., J.B., S.A., J.P., G.G., T.K., M.L., N.N., O.L., E.B.E., R.M., S.R., H.S., S.S., F.T., and R.B-R collected the data. E.D., J.B., J.P., E.B.E., R.M., S.R., H.S., F.T., and R.B-R contributed data or analysis tools. E.D., J.B., G.G., T.K., M.L., N.N., A.R., L.B., E.B.E., R.M., F.T., and R.B-R performed the analysis. S.A., S.L., D.M., A.P., S.S., F.T., and R.B-R contributed intellectual input/interpretation. S.A., F.T., and R.B-R wrote the paper.

## Ethics approval and consent to participate

Informed consent was obtained for all patients. The study was in strict compliance with all institutional ethical regulations.

## Supplementary Figures

**Supplementary Figure 1.**
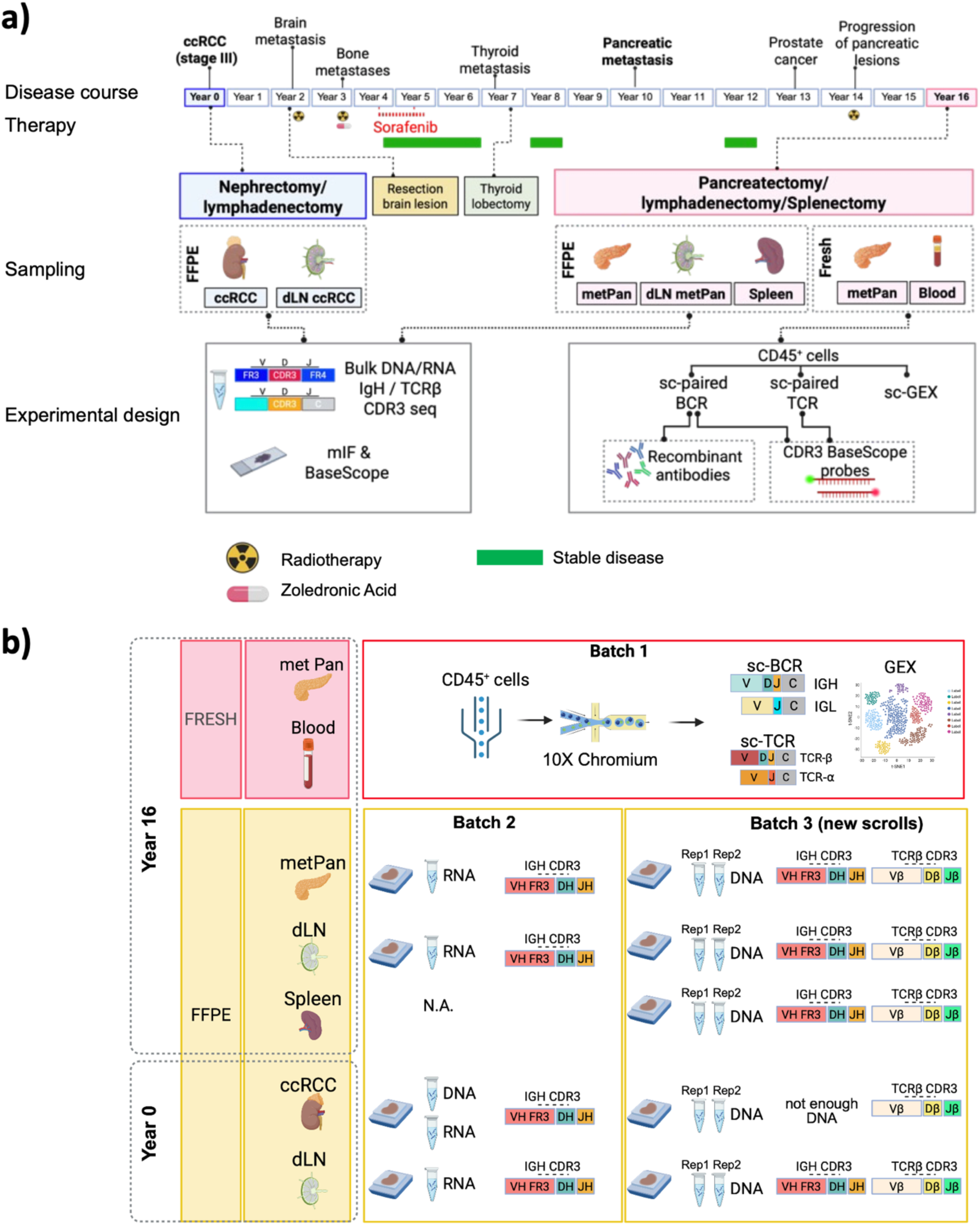
**a)** Clinical history and experimental set up. **b)** Detailed schematic of BCR and TCR repertoire analysis across tissue sites, highlighting tissue batches, DNA and/or RNA input, and PCR amplicons. The detection of shared B cell CDR3 sequences across batches and methodologies strongly supports the robustness and reproducibility of data, providing compelling evidence of B cell clonal persistence, and effectively ruling out PCR contamination artifacts.

**Supplementary Figure 2.**
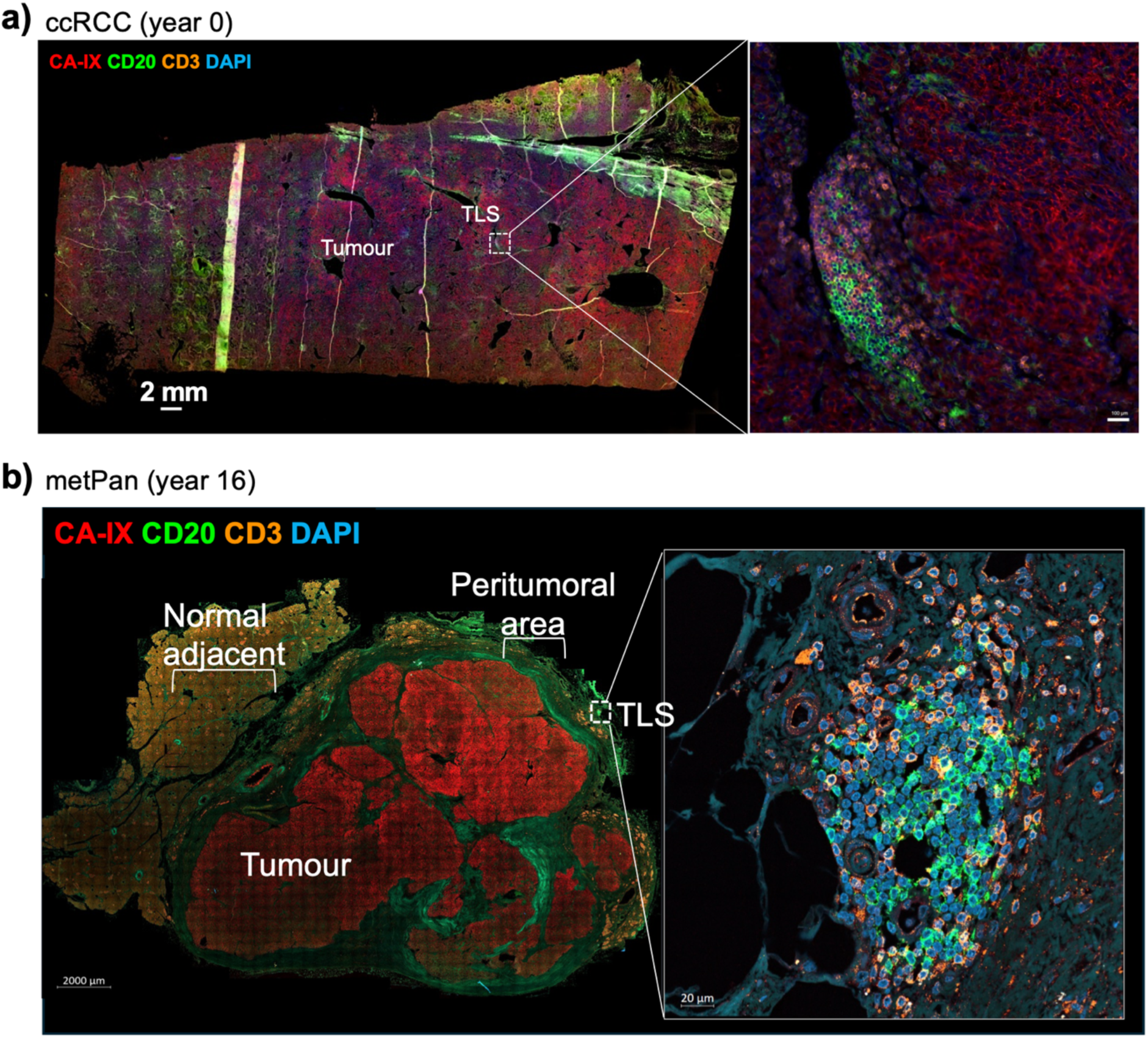
TLS proximity to CA-IX expressing tumour cells in ccRCC and metPan. Representative mIF images of the primary ccRCC **(a)** and metPan **(b)** FFPE tissue samples stained with anti-CA-IX combined with anti-CD20 and anti-CD3 antibodies; DAPI for nuclei. **a)** In ccRCC primary tumour, TLS are in close proximity to CA-IX positive cells. The TLS magnification shows co-localization of CD20^+^ B cells with CD3^+^ T cells (scale bar= 100µM). **b**) In metPan TLS are confined in the peritumoral region. No TLSs were detected within the CA-IX positive tumour nodes or in the normal adjacent area. The TLS magnification shows co-localisation of CD20^+^ B cells with CD3^+^ T cells (same image as in Figure 3c).

**Supplementary Figure 3.**
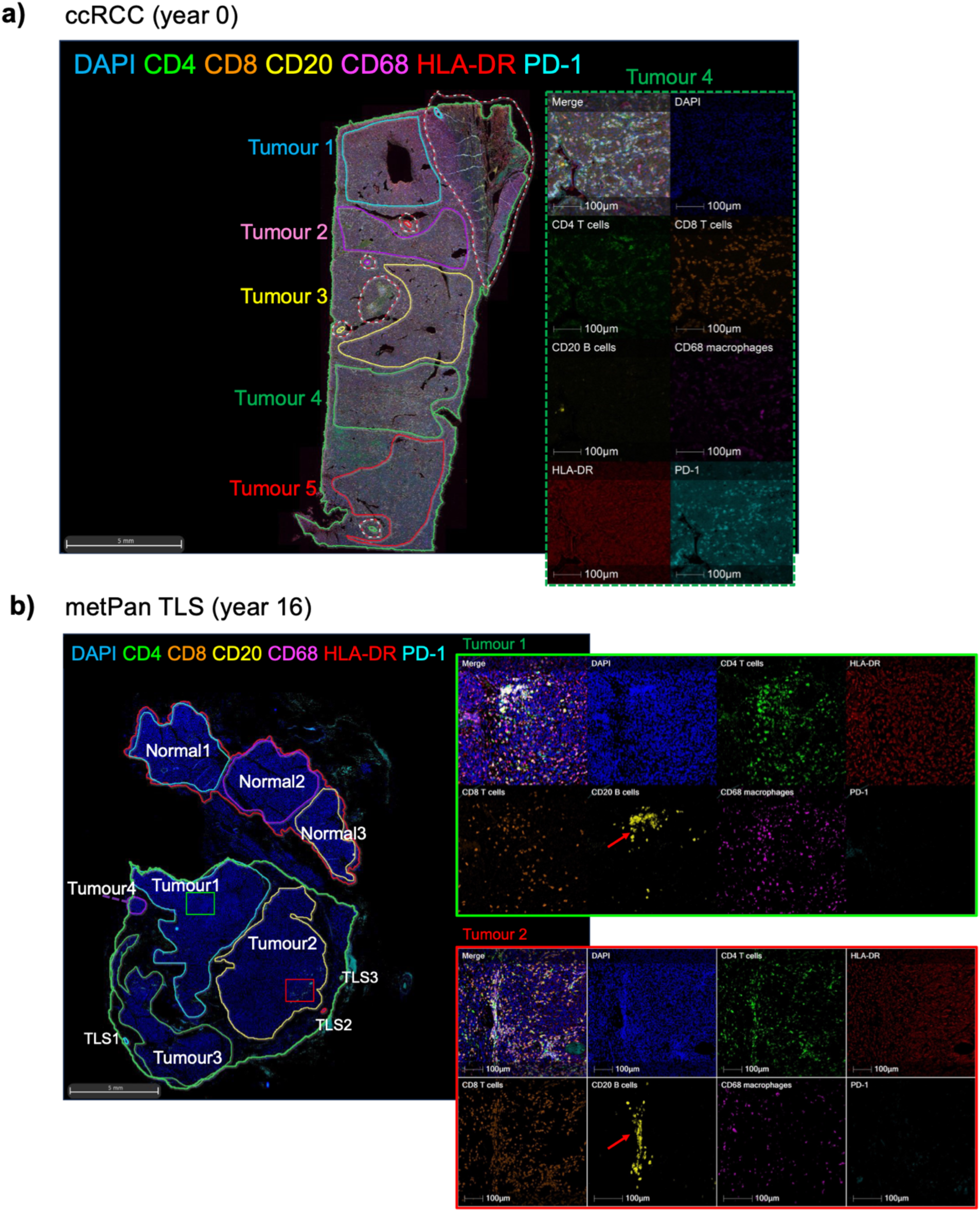
Representative mIF images of ccRCC and metPan with tumour ROI and TLS annotations (related to Figure 1). In both metPan and ccRCC, tumour ROIs have been selected to exclude TLS and to represent the variability of immune cell infiltration throughout tissue sections. **a)** Representative mIF image of ccRCC showing 5 annotated tumour ROIs (Tumour 1-5) throughout the TME. The single channel shows the distribution of immune cells within tumour ROI 4. **b)** Representative mIF image of metPan section with TLS, tumour and adjacent normal ROI annotations. The single channel image shows two ROIs, from tumour 1 and tumour 2. While most immune cells are found both clustered and scattered throughout the TME, B cells are detected exclusively within clusters (red arrows).

**Supplementary Figure 4.**
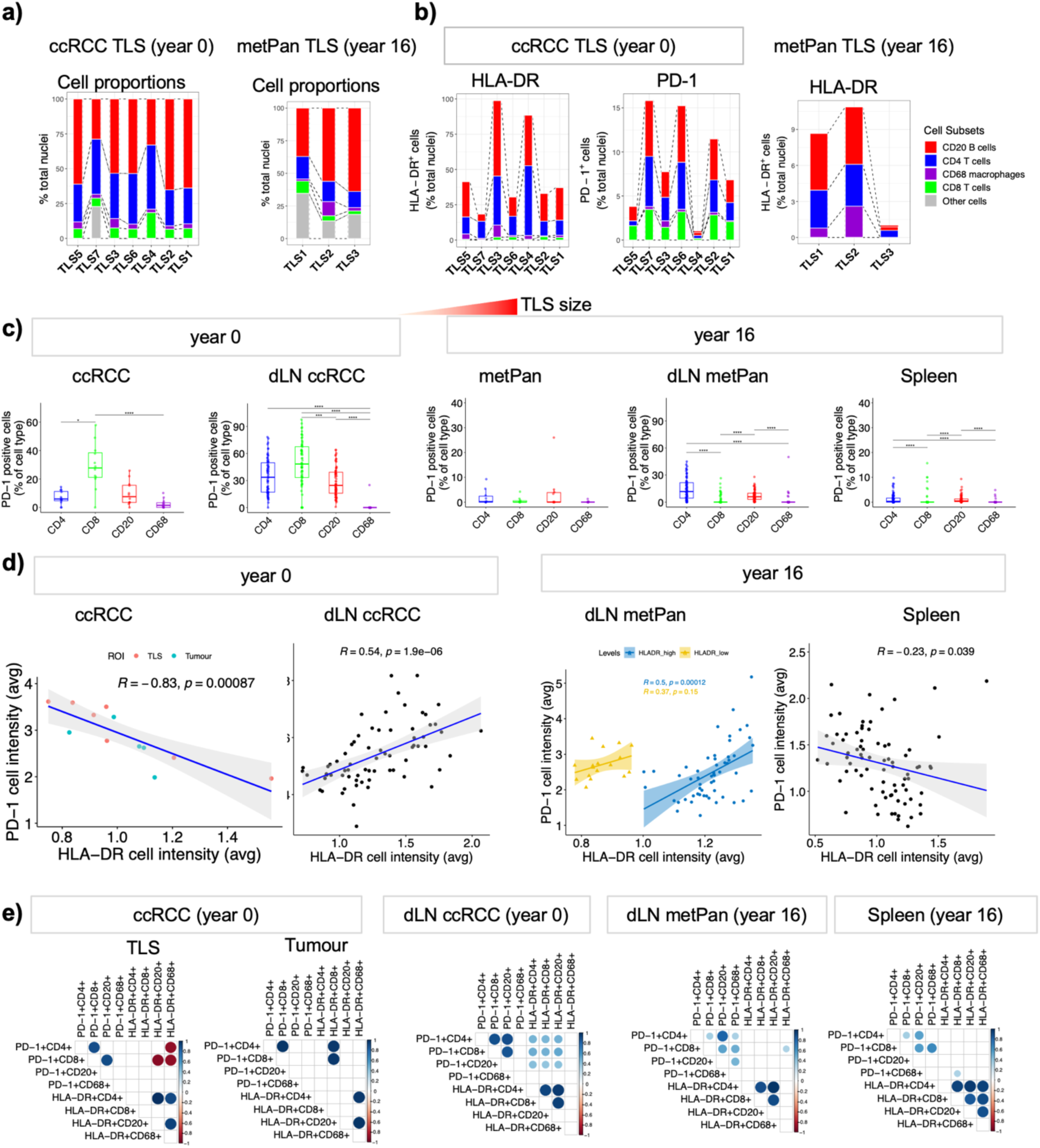
PD-1 and HLA-DR expression profiles across TLS and dLN GCs (related to Figure 1). a) Distribution of PD-1^+^ cells across tumour sites, dLN, and the spleen at year 0 and 16. Box plots showing the frequency of PD-1^+^ cells (as % of cell type). PD-1^+^ cells are particularly enriched among CD8 T cells in ccRCC and its dLN. b) Cellular composition and functional status of TLS based on size. Alluvial plots showing the cellular composition, the proportion of PD-1^+^ and HLA-DR^+^ cells (as % of nuclei) across TLS ordered by size in ccRCC and metPan. c) Correlation between PD-1 and HLA-DR expression within ccRCC, dLNs, and the spleen. Scatter plots with linear regression of PD-1 and HLA-DR average cell intensity across sites. In ccRCC, dots represent TLS and tumour ROIs. In dLNs and spleen, dots represent GCs as annotated on images in Figure 1. In metPan dLN, HLA-DR intensity was separated into high >1, and low <1 intensity for a better visualization of correlation. (Pearson correlation coefficient and p-value are shown). d) Correlation between the percentage of PD-1 and HLA-DR positive cells (as percentage of cell subsets) within ccRCC (tumour and TLS), and GCs of the dLN the spleen. The color and the values indicate the Pearson correlation coefficient (significance ≤0.01).

**Supplementary Figure 5.**
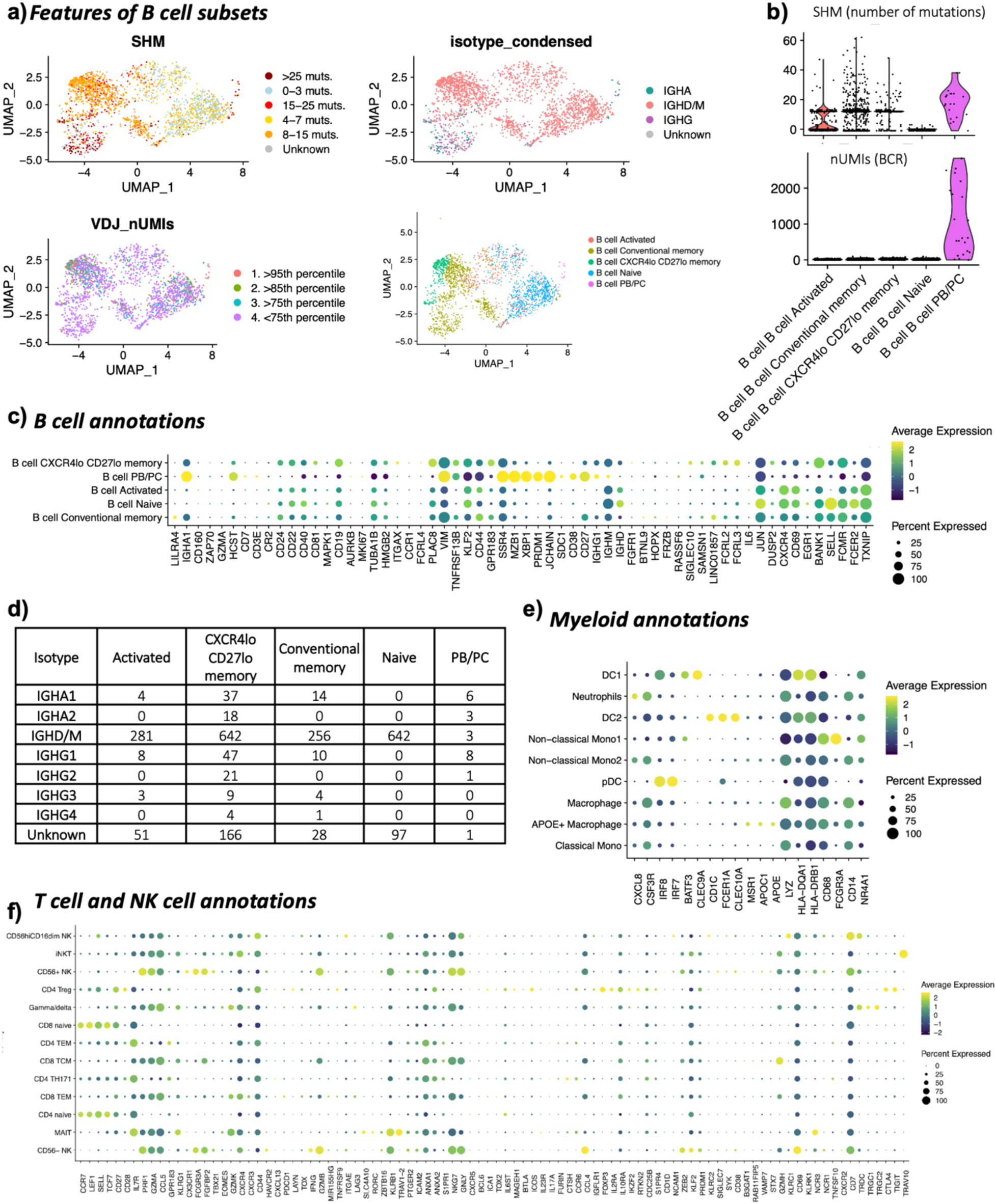
Single cell annotation and clustering of blood and metPan CD45^+^ cells (related to Figure 2). a) UMAP plots of B cell clusters showing SMH, isotype, number of BCR heavy chain UMIs, grouped into percentile groups, B cell subsets; b) Violin plots of the number of somatic hypermutations SHM (above) and number of BCR heavy chain reads (UMIs) per B cell subset; c) Gene expression profiles of B cell subpopulations of the key differentially expressed genes. d) Isotype usage count (number of cells) per B cell subset. e) Gene expression profiles of myeloid cell subpopulations of the key differentially expressed genes. f) Gene expression profiles of T and NK cell subpopulations of the key differentially expressed genes.

**Supplementary Figure 6.**
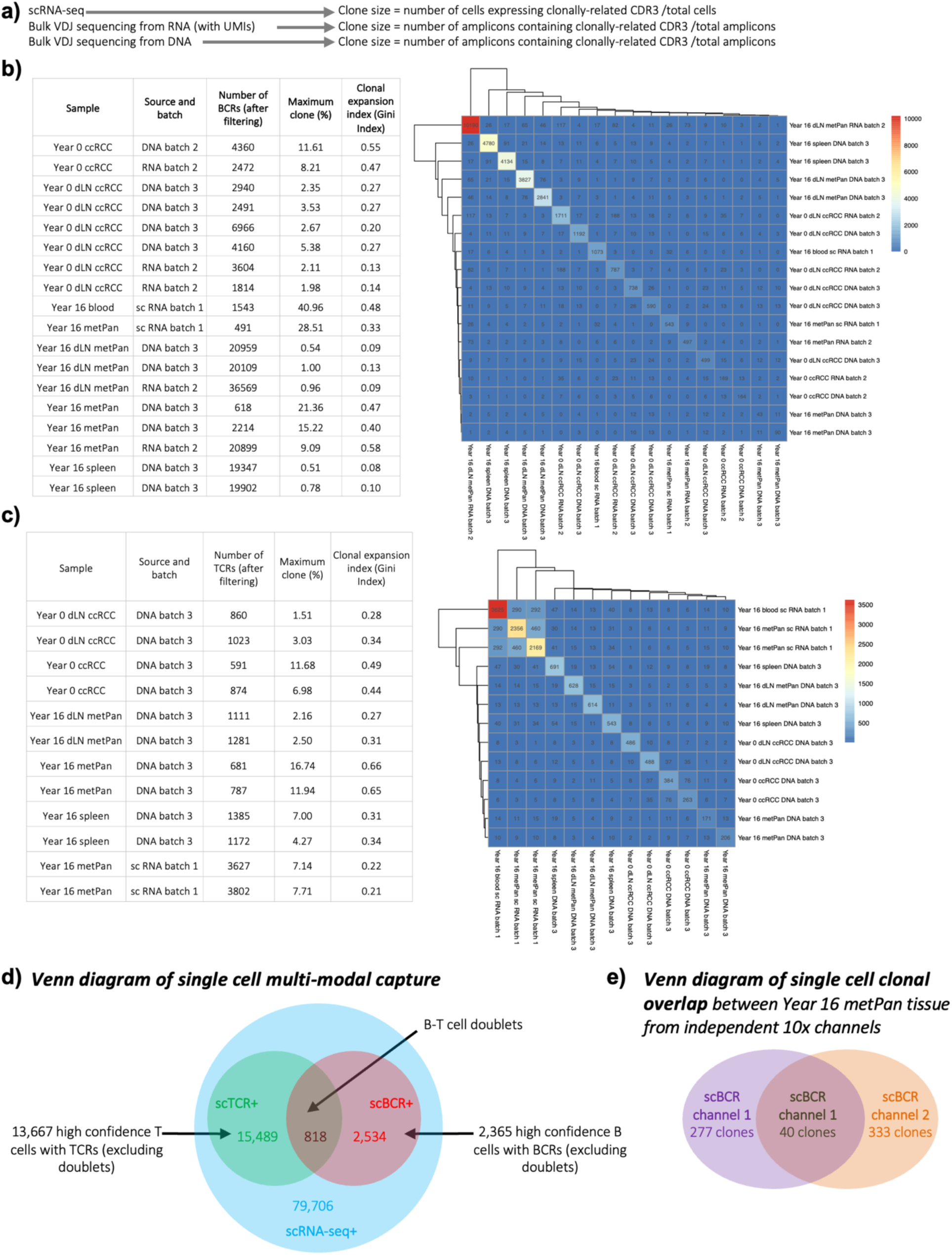
BCR and TCR VDJ sequencing summary. a) Schematic of clone definitions between different sequencing methods from different sample sources. b) Summary table of BCR capture, maximum clone size and clonal expansion Index across all samples (left) and heatmap of the counts of overlapping clones between samples. c) Summary table of TCR capture, maximum clone size and clonal expansion Index across all samples (left) and heatmap of the counts of overlapping clones between samples. d) Venn diagram of single cell multi-modal capture across Year 16 metPan tissue and blood samples. e) Venn diagram of single cell clonal overlap between Year 16 metPan tissue from independent 10x channels.

**Supplementary Figure 7.**
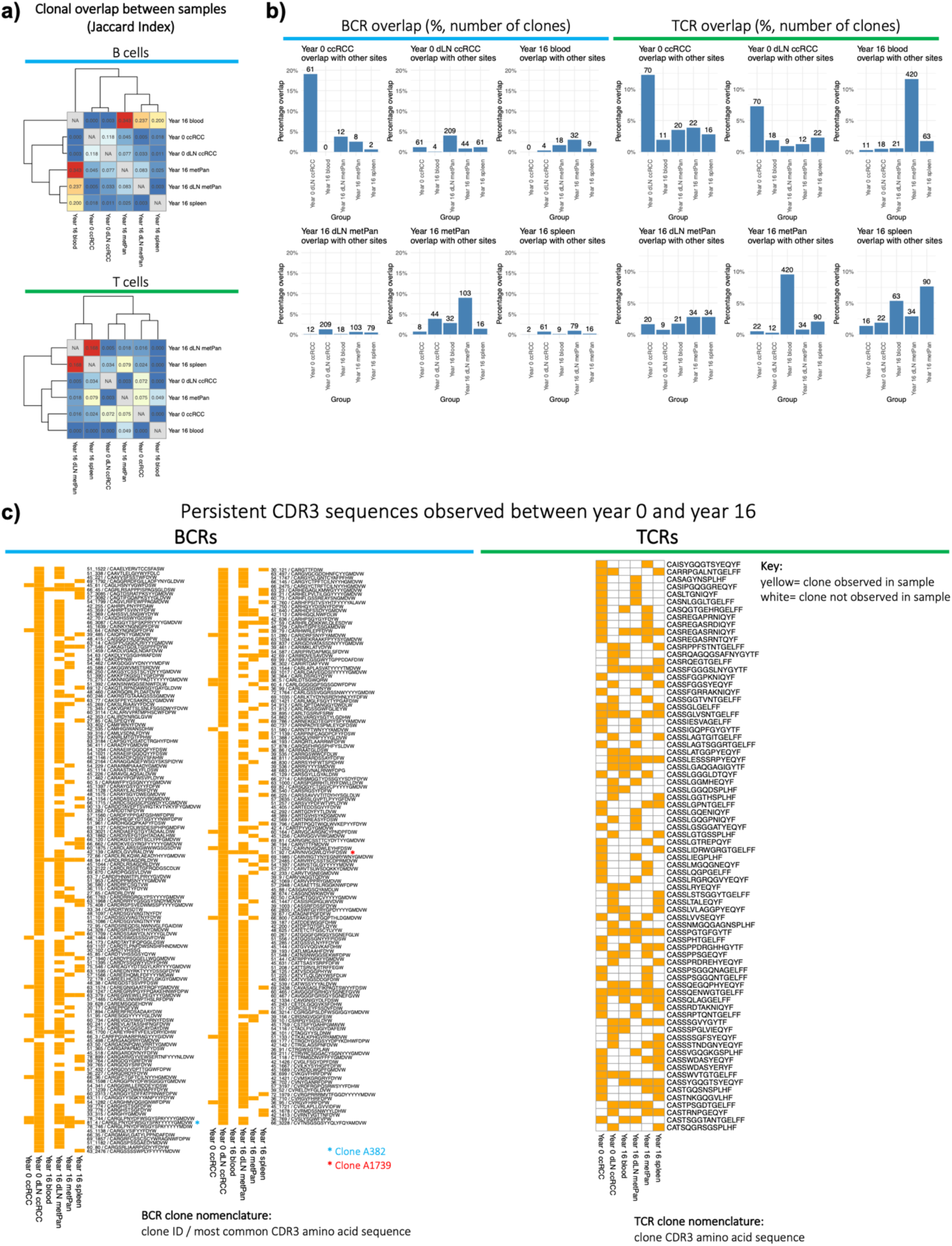
B and T cell clonal sharing and persistence. a) Clonal overlap (Jaccard Index) between sites and time points for (left) BCRs and (right) TCRs. b) Bar plots illustrating clonal sharing of BCRs (left) and TCRs (right) between different anatomical sites. The height of each bar represents the percentage of shared clones, with the total number of shared clones indicated above each bar. b) Heatmap showing the persistence of BCR and TCR clones between diagnosis (Year 0) and the 16-year follow-up (Year 16) across all samples. Yellow boxes indicate the presence of a specific clone in a given sample. Blue and red asterixis indicate the clones in Figure 2e.

**Supplementary Figure 8.**
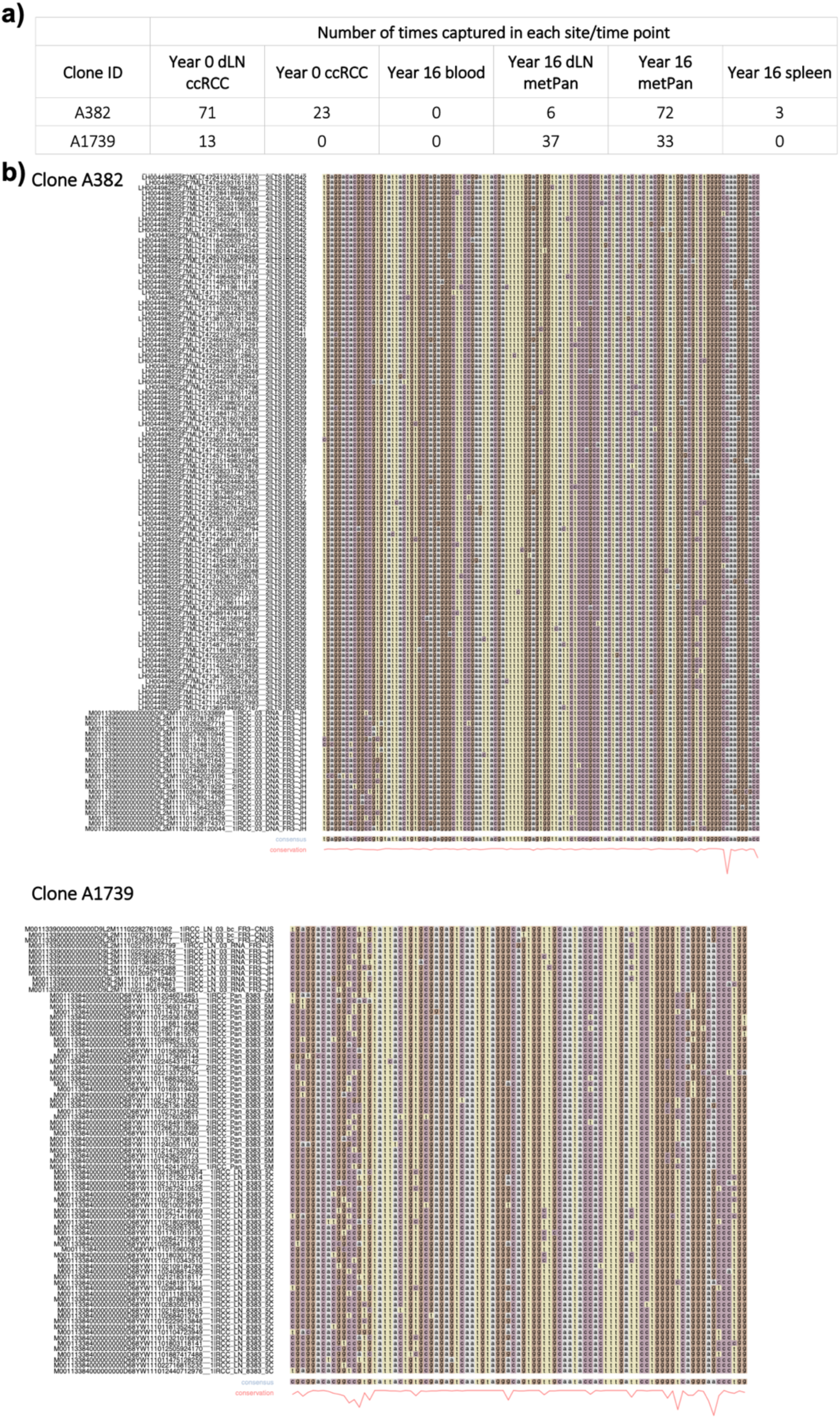
Expanded persistent BCR clonal information from Figure 2e. a) Clonal counts for persistent BCR clones across all samples. b) Heavy chain sequence alignments for the persistent BCR clones. The conservation score for each alignment is shown beneath the sequences.

**Supplementary Figure 9.**
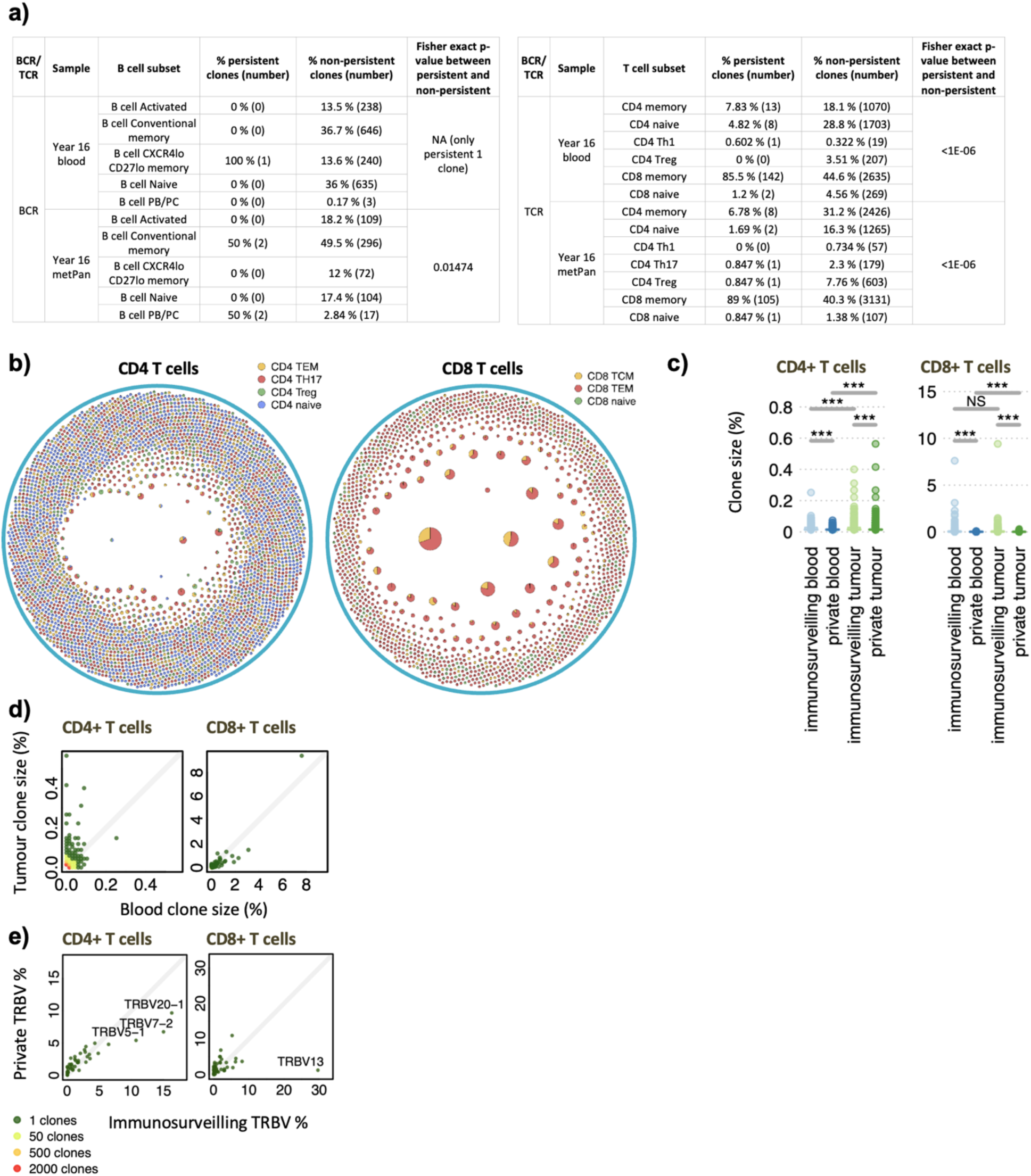
Features of persistent clones and TCR repertoire data. a) Tables showing the number and percentage of persistent versus non-persistent cells for each B cell (left) and T cell (right) subset, along with Fisher exact p-values per sample. b) VDJ clonality network plots for CD4 (left) and CD8 (right) T cells. Each node represents a unique VDJ sequence, with its size indicating the number of cells. Each node is a pie chart showing the proportion of cells belonging to each T cell subset. c) Box plots of the relative single-cell clone sizes between clones that are shared between year 16 blood and year 16 metPan samples, separated by CD4 and CD8 phenotypes. d) Scatter plot showing the clone sizes of T cell clones found in both Year 16 blood and Year 16 pancreatic metastasis samples. e) Bar plot of the TRV gene usages for T cell clones found in both the Year 16 blood and Year 16 pancreatic metastasis samples.

**Supplementary Figure 10.**
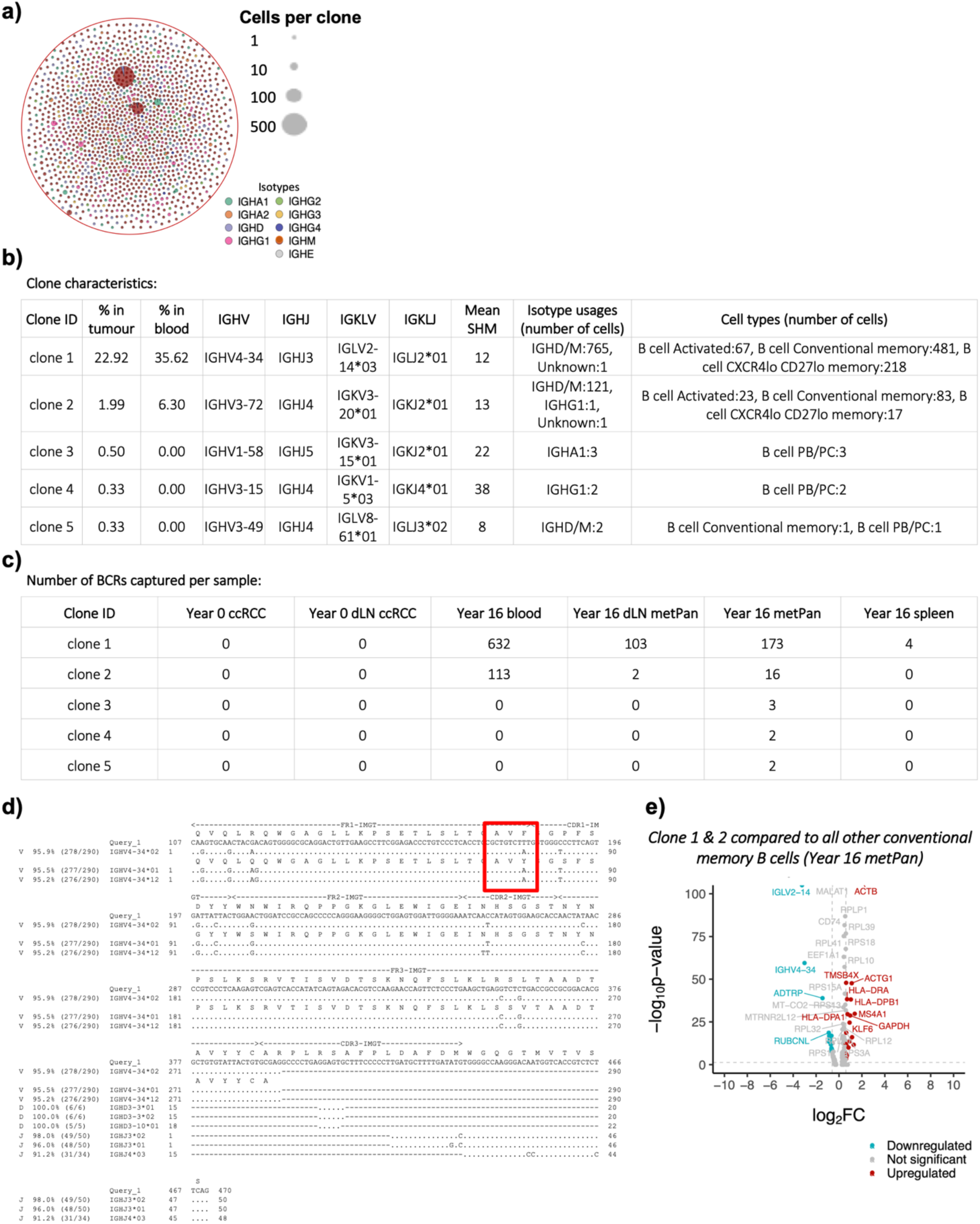
BCR clonality network and features of the selected B cell clones for antibody expression. a) VDJ clonality network plots for B cells, where each node represents a unique VDJ sequence, with its size indicating the number of cells. Pie charts within each node show the proportion of cells expressing each isotype. Selected clones for monoclonal antibody production are indicated. b) Key characteristics of the selected B cell clones used for monoclonal antibody production. c) Number of BCRs captured per sample for selected clones from which monoclonal antibody production was performed. d) Ig-Blast annotation and alignment of the heavy chain sequence for Clone 1. The region previously linked to autoreactivity is highlighted in red. e) Volcano plot from differential gene expression (DGE) analysis comparing Clone 1 and Clone 2 to other conventional memory B cells from the Year 16 pancreatic metastasis sample.

**Supplementary Figure 11.**
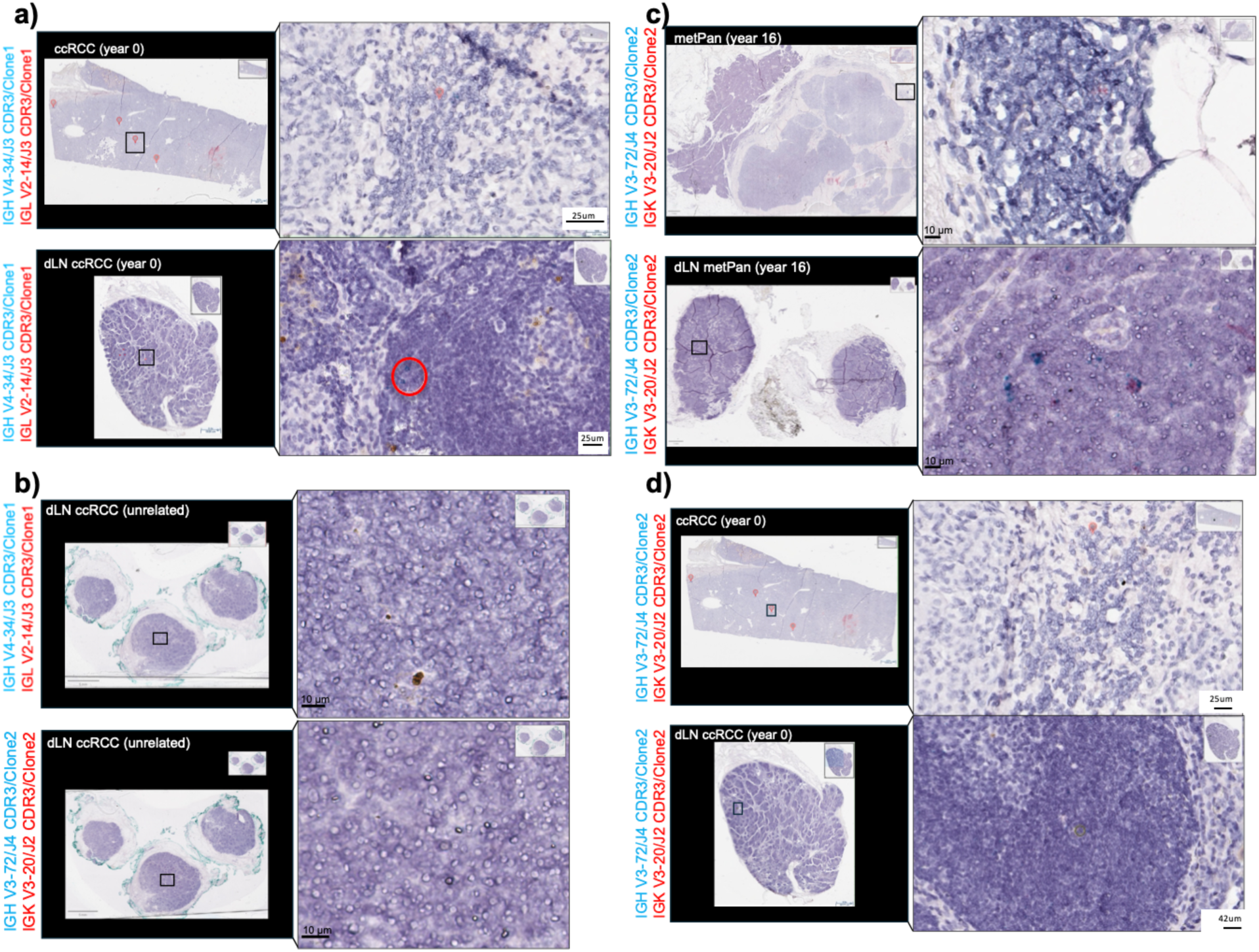
BaseScope assay to detect clone 1 and clone 2 BCR transcripts across tumour sites and draining lymph nodes (dLN) (related to Figure 3). The Basescope assay reveals the spatial distribution of IgH and IgL CDR3s as blue dots and red dots, respectively. Clone 1 and clone 2 V(D)J genes are annotated on the left-hand side of the images. a) Representative images of ccRCC (top) and ccRCC dLN (down) with B cell Clone 1 probes (IgH + IgL). The red pins indicate small immune clusters within the TME. Very few instances of clone 1 B cells were detected within the ccRCC dLN (red circle). b) Representative image of an unrelated ccRCC dLN. The specificity of B cell clone 1/clone 2 CDR3 probes was tested against the dLN from an unrelated ccRCC dLN (tissue obtained in the same year as metPan/metPan dLN/spleen). No signal was detected on the unrelated ccRCC dLN, indicative of probe specificity. c) Representative images of metPan (top) and metPan dLN (down) with B cell Clone 2 probes (IgH + IgL). Very few dots of clone 2 B cell CDR3 mRNA were detected in the metPan TLS and the pancreatic dLN. The red pin in metPan indicates the peritumoral TLS as in Figure 3b. The red circles in metPan dLN indicate a few examples of clone 2 CDR3 transcript detection (IgH + IgL). d) Representative images of ccRCC (top) and ccRCC dLN (down) with B cell Clone 2 probes (IgH + IgL). The red pins indicate small immune clusters within the TME. No instances of clone 2 B cells were detected in these sections. Slides acquired with the Leica Aperio system, 40X objective.

**Supplementary Figure 12.**
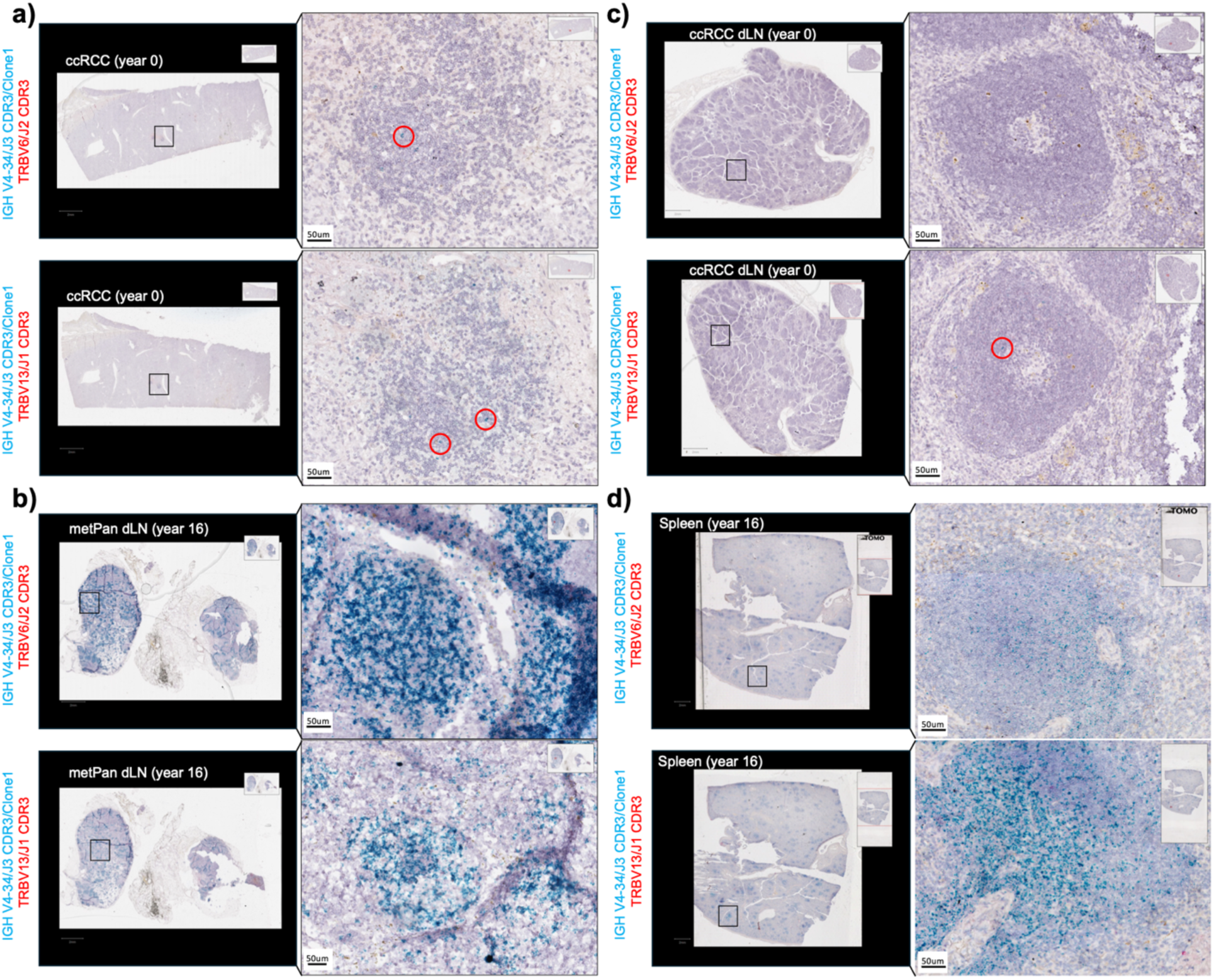
Mapping the spatial co-localisation of Clone 1 B cells (VH4-34) with two clonally expanded T cell clones (Clone 1:TRBV6; Clone 2: TRBV13). Clone 1 IGH CDR3 (VH4-34) probe (same as in Figure 3b) was combined with probes targeting the TCRβ CDR3 of Clone 1 (TRB-V6) and Clone 2 (TRB-V13) T cells. Red dots indicate either TRB-V6 or TRB-V13 CDR3 transcripts, and blue dots indicate Clone 1 IGH CDR3 transcripts. The combination of V(D)J genes is reported on the left-hand side of the images. a) Representative image of ccRCC stained with Clone 1 IGH CDR3 probe combined with Clone 1 TRBV6 (top) and Clone 2 (TRBV13) (down) T cell probes. A few instances of Clone 1 B cells were detected within the largest intratumoral TLS in ccRCC (red circles). No instances of TRBV6 and TRBV13 T cells were found. b) Representative image of ccRCC dLN stained with Clone 1 IGH CDR3 probe combined with Clone 1 TRBV6 (top) and Clone 2 (TRBV13) (down) T cell probes. A few instances of Clone 1 B cells were detected (red circle). No instances of TRBV6 and TRBV13 T cells were found. c) Representative image of metPan dLN stained with Clone 1 IGH CDR3 probe combined with Clone 1 TRBV6 (top) and Clone 2 (TRBV13) (down) T cell probes. As in Figure 3b, Clone 1 B cells populated most of the follicles and GCs. No instances of TRBV6 and TRBV13 T cells were found. d) Representative image of the spleen stained with Clone 1 IGH CDR3 probe combined with Clone 1 TRBV6 (top) and Clone 2 (TRBV13) (down) T cell probes. Clone 1 B cells were detected within splenic GCs, though with lower intensity compared to metPan dLN. No instances of TRBV6 and TRBV13 T cells were found.

**Supplementary Figure 13.**
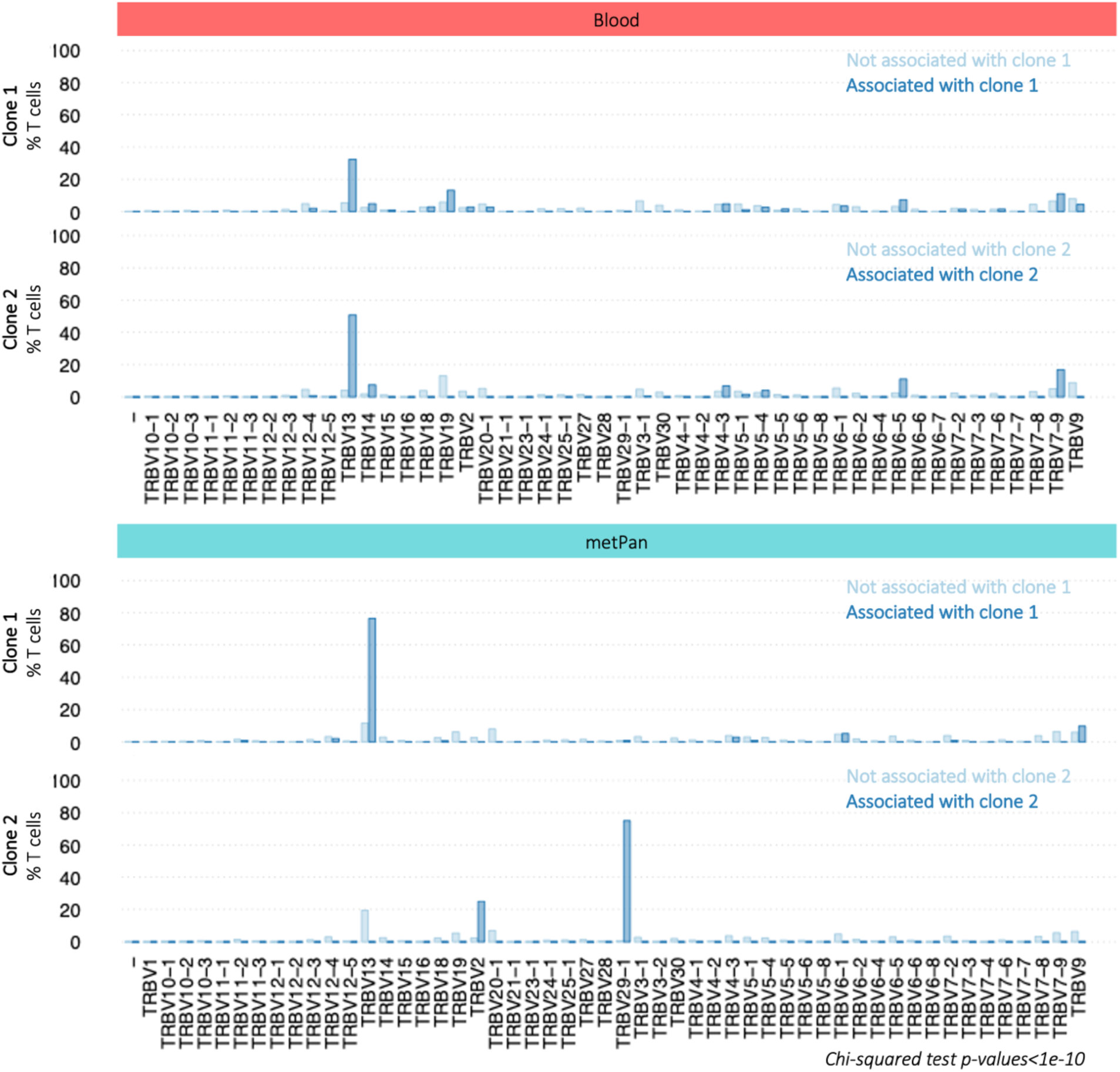
Bar plots showing the proportion of T cell subsets that are clonally related to T cells found in doublets with Clone 1 B cells (top) or Clone 2 (bottom) in Year 16 blood and Year 16 metPan samples. This is compared to the proportion of T cells that are not doublet-related (light blue).

**Supplementary Figure 14.**
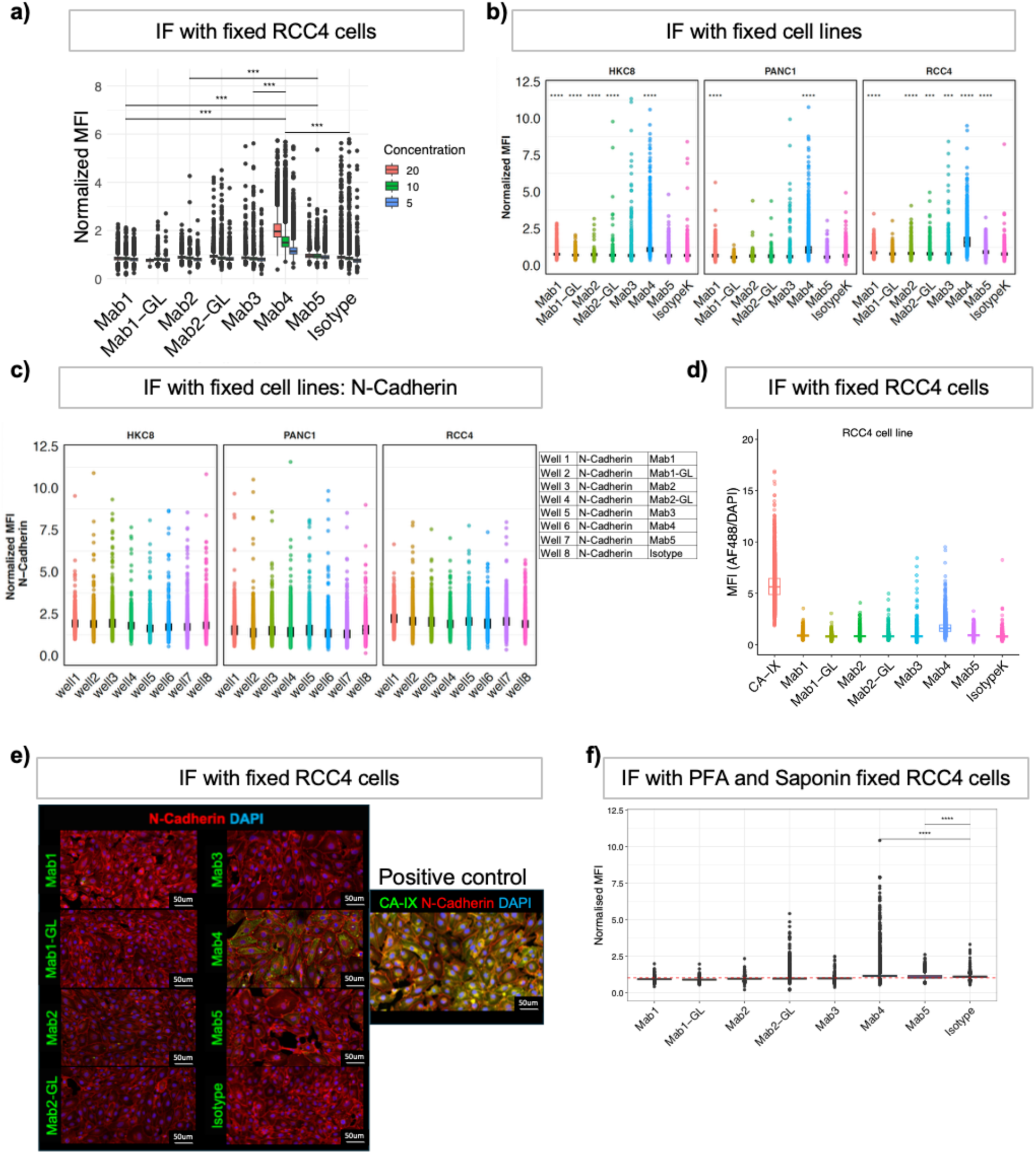
Immunofluorescence staining of epithelial cell lines using monoclonal and control antibodies (related to Figure 4). a) Titration of the monoclonal antibody concentration [µg/mL] for IF staining of RCC4 cells. Boxplots representing the mean fluorescence intensity (MFI) normalised to DAPI of each antibody at different concentrations. Each dot represents a single object (cell) obtained through segmentation and annotation with the Arivis Software implementing a Cellpose pipeline for cellular segmentation. Statistics was performed by Kruskal-Wallis test followed by post-hoc Dunn test (***= p value < 0.001). b) Bar plots showing Dunn’s Test Z scores for the comparison across antibodies, highlighting that Mab4 and Mab5 antibodies stand out as significantly different from most other antibodies and the isotype control. c) Boxplots showing normalised MFI of cell lines stained with monoclonal antibodies and the isotype control. Statistics was performed by one-sided Wilcoxon rank-sum test for each monoclonal antibody against the isotype ctrl., significant p values are shown (****, p ≤ 0.0001). d) Boxplots showing normalised MFI of the anti-N-Cadherin staining that was performed in combination with monoclonal antibodies across cell lines. e) Representative IF images of RCC4 cell line stained with monoclonal antibodies in combination with anti-N-Cadherin, and with anti-CA-IX in combination with N-Cadherin. DAPI for nuclei. The images show a higher signal with Mab4 as compared to other monoclonal antibodies. The image with anti-CA-IX antibody has been used as a positive control to assess intracellular staining. Images were acquired with the Olympus Spinning Disk, 20X magnification. f) Boxplots showing normalised MFI of RCC4 cells stained with anti-CA-IX (Carbonic Anhydrase 9) antibody in comparison to monoclonal antibodies. g) Boxplots of normalised MFI of RCC4 cells stained with monoclonal antibodies and isotype control upon PFA 4% and saponin 0.1% fixation and permeabilisation. The data confirm a higher binding of Mab4 compared to other antibodies. Statistics was performed using Wilcoxon rank-sum test for each group against the reference group (Isotype) (****p<0.001).

**Supplementary Figure 15 (related to Figure 5).**
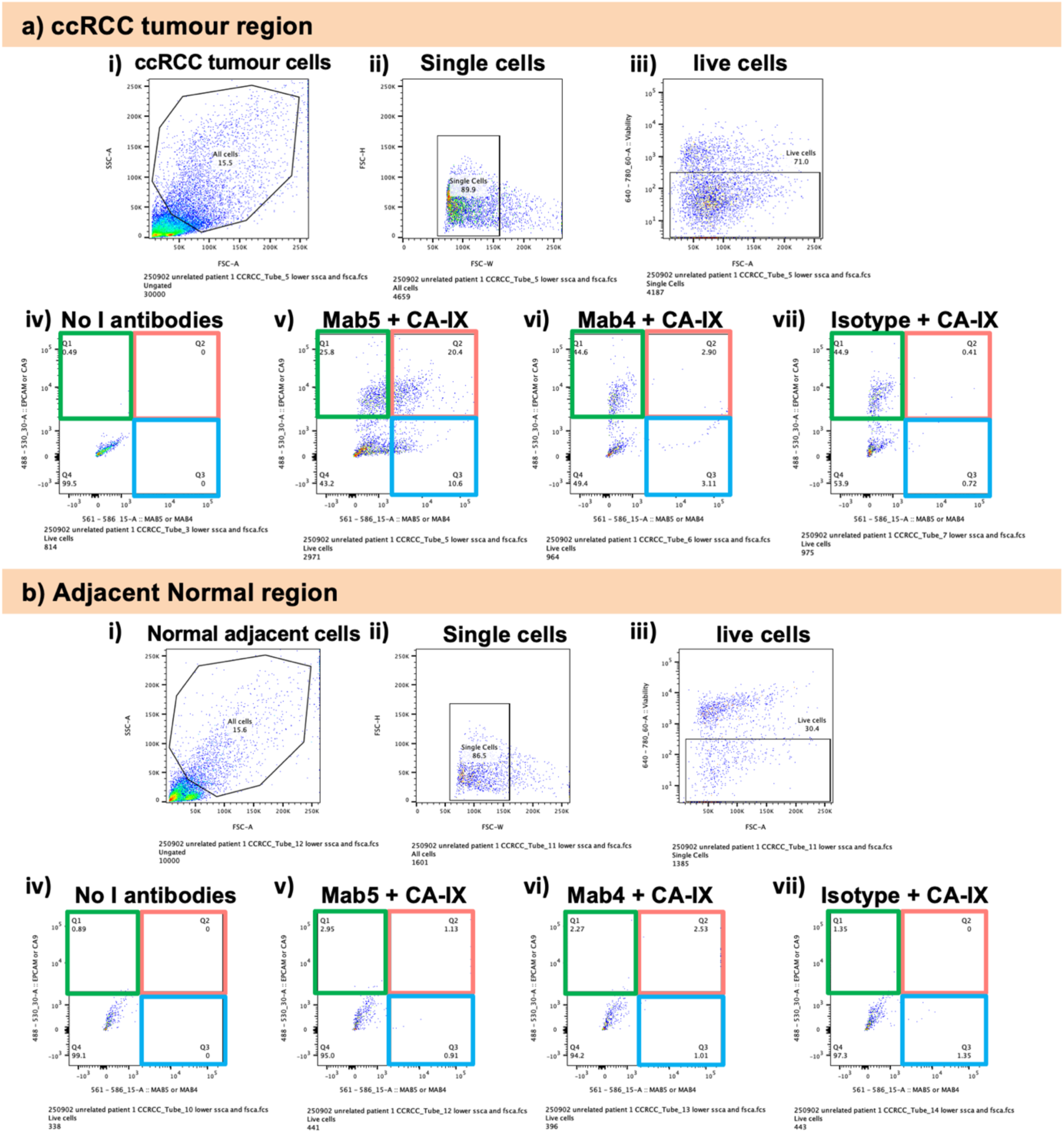
Flow cytometry analysis (FACS) of (a) dissociated ccRCC tumour and (b) normal adjacent tissue from an unrelated patient, using the monoclonal Mab4 and Mab5 antibodies in combination with the anti-CA-IX antibody and controls. The gating strategy for both ccRCC and adjacent normal cells is shown: i) tumour/normal cells were selected based on forward scatter (FSC) and side scatter (SSC) parameters; ii) doublets and aggregates were excluded using FSC-H versus FSC-W; iii) live cells were identified using LIVE/DEAD™ Fixable Near-IR Dead Cell Stain; iv) A negative control without primary antibodies was included to determine background signal; v) cells were stained with anti-CA-IX (Carbonic Anhydrase 9) and Mab5; vi) cells were stained with anti-CA-IX and Mab4; vii) cells were stained with anti-CA-IX and a mouse isotype control antibody.

**Supplementary Figure 16.**
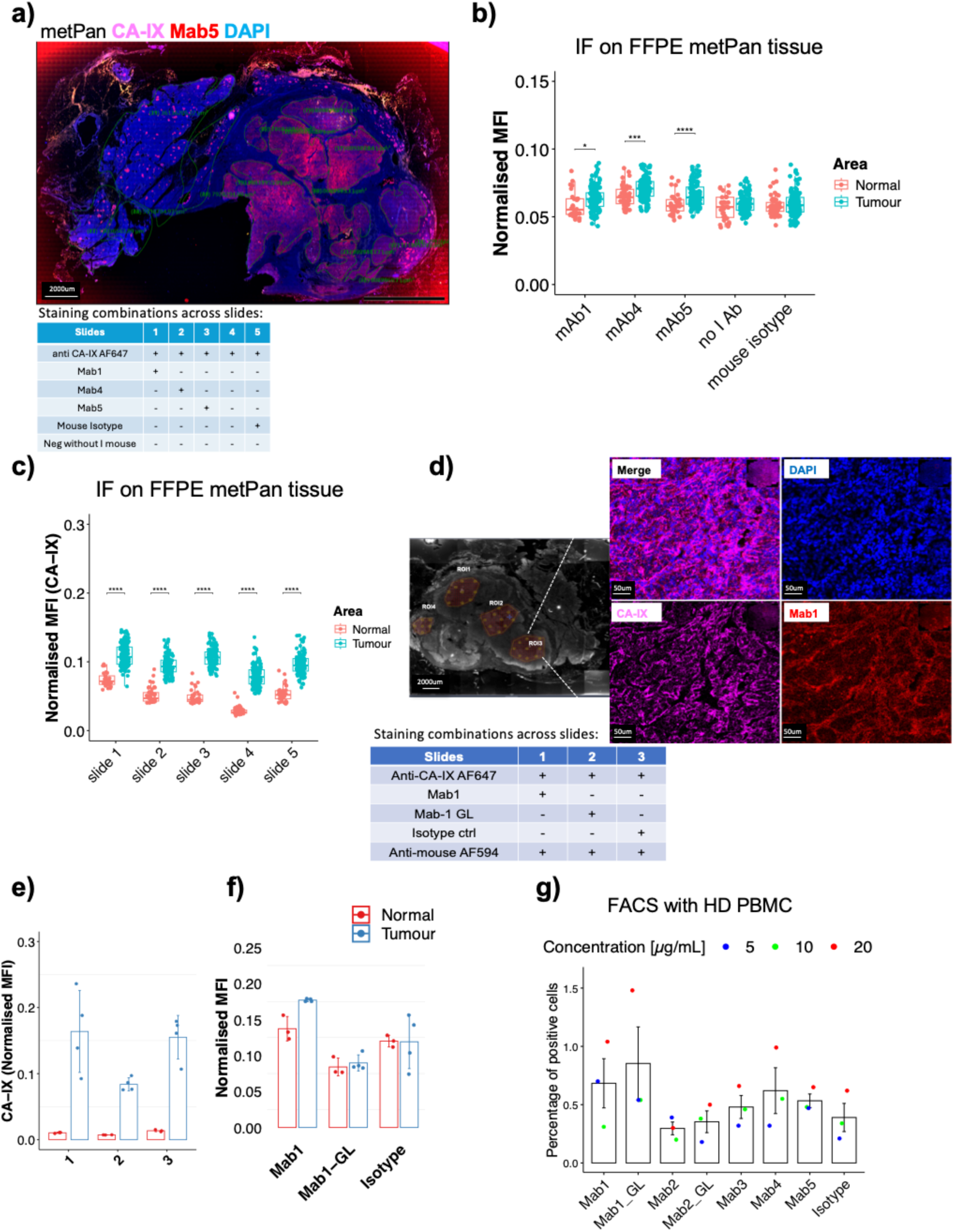
Reactivity of the monoclonal antibodies (Mabs) against the patient FFPE metPan tissue and healthy donor PBMC. a) Representative immunofluorescent image of a metPan healthy donor PBMC and FFPE section stained with the monoclonal Mab5 combined with the anti-CA-IX AF647 (Carbonic Anhydrase 9) antibody. Several ROIs have been selected within the tumour and the adjacent normal areas. Highly autofluorescent histological structures were excluded from the analysis. The table shows the schematic of each slide staining, with antibody combination. b) Grouped boxplots of normalised MFI for Mab1, Mab4, Mab5, mouse isotype control, and negative control without the mouse primary antibody, in normal and tumour areas within metPan. Statistics was performed by Mann Whitney U test between Normal and Tumour areas, significant p-values are shown (*, p =0.012; ***, p = 0.00015; ****, p = 0.00007). c) Grouped boxplots showing normalised MFI of CA-IX staining between normal and tumour areas across each slide of metPan. Statistics shows p-values of Mann Whitney U test between Normal and Tumour areas (****, p. adj <0.0001). d) Representative immunofluorescent image of a metPan FFPE section stained with the monoclonal Mab1 combined with anti-CA-IX AF647 (Carbonic Anhydrase 9) antibody. The image on the left-hand side shows whole section scanning with annotated ROI within the tumour area. The magnifications on the left-hand side shows the staining and the single channels. The table shows the schematic of each slide staining. e) Grouped bar plots showing normalised MFI of CA-IX staining between normal and tumour ROI across metPan FFPE slides. f) Grouped bar plots of normalised MFI of Mab1-, Mab1-GL, and isotype control in normal and tumour areas across metPan FFPE slides. Statistics were performed by Wilcoxon pairwise comparison with Bonferroni correction across monoclonal antibodies (p=ns) and by Wilcoxon pairwise test between Normal and Tumour ROI. Each dot on the bar plot represents a large ROI across the tissue. Representative immunofluorescent image of a metPan FFPE section stained with the mouse monoclonal Mab1, Mab4, Mab5 combined with anti-CA-IX AF647 (Carbonic Anhydrase 9) antibody, compared to controls. g) Poly-reactivity of the monoclonal antibodies was assessed by FACS using healthy donor (HD) PBMC. Bar plots show the percentage of positive cells across three antibody concentrations. While the comparison of positive cells across antibodies did not reach statistical significance, a trend of higher reactivity was observed for Mab1-GL (Kruskal-left-hand chi-squared = 8.4173, df = 7, p-value = 0.2972).

**Supplementary Figure 17.**
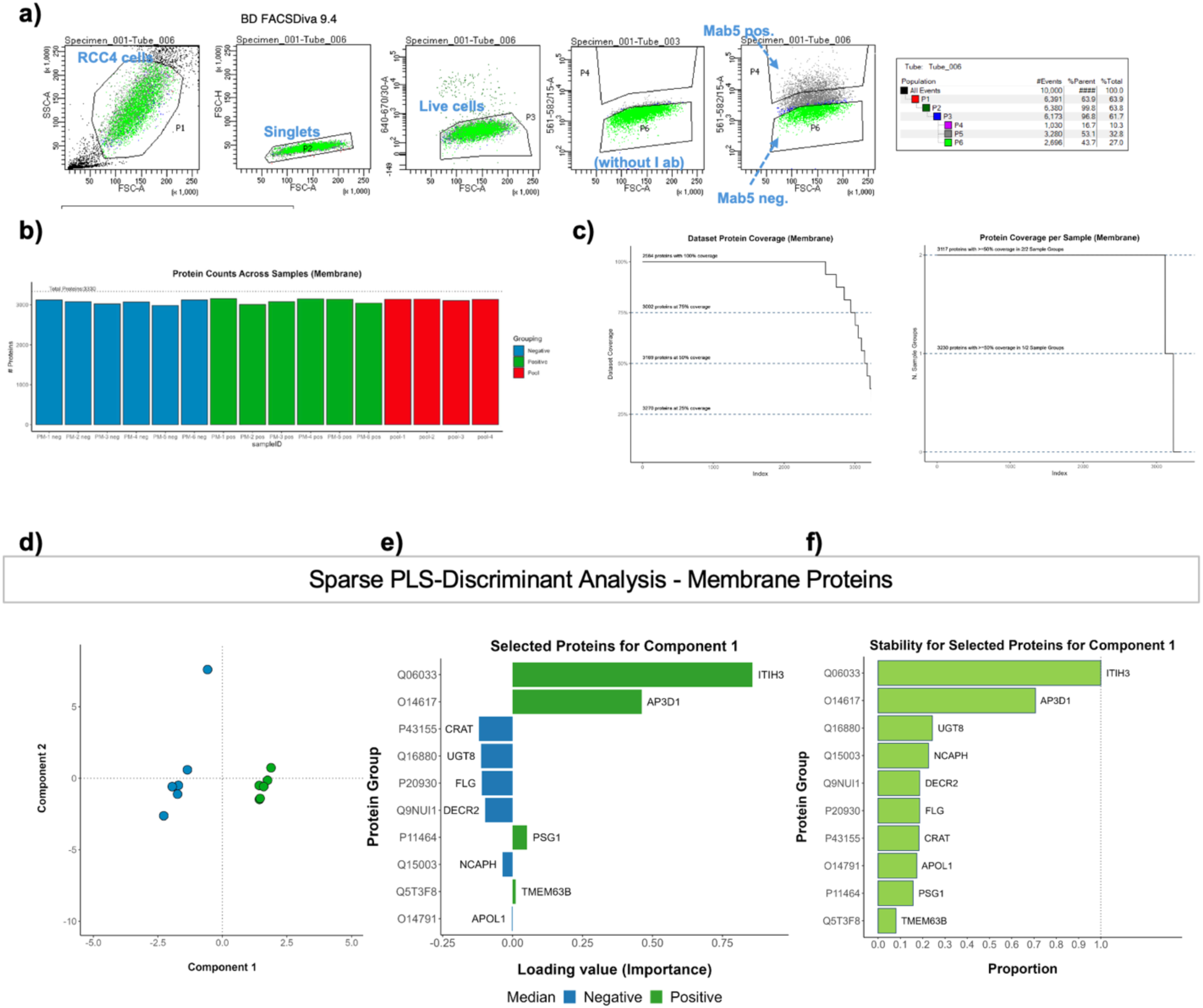
Mass spectrometry of plasma membrane proteins to identify Mab5 antigenic targets on RCC4 cells (related to Figure 4). a) Schematic of flow cytometry and cell sorting (FACS) of Mab5 positive and Mab5 negative RCC4 cells. The plots show the gating of singlets, live cells, and Mab5 positive and negative groups based on a negative control threshold (without Mab5 antibody). b) Dataset metrics with plasma membrane count in Mab5-positive, Mab5-negative and pooled groups. Membrane proteins are defined as proteins that are present in the AFTM database, present in the MatrisomeDB database, or annotated with the membrane GO term (GO:0016020). c) There were 2584 membrane protein groups that were quantified across all samples. There were 3220 membrane protein groups that were quantified in at least 50% of at least one sample group (negative and positive only, excluding pool group). Including the protein groups with 50% coverage in at least one sample group allows for inclusion of proteins completely missing in one sample group while maintaining statistical power. d) Sparse PLS-Discriminant analysis of membrane proteins using normalised log2 intensities with positive and negative as target classes. Clustering of positive and negative groups with component 1. e) Bar plot showing 10 proteins selected for component 1 and ordered by loading value. f) Bar plot showing stability scores for proteins selected in component 1.

## Supplementary Tables

**Supplementary Table 1.** Statistical significance of the comparison of cellular composition across SLO (dLNs and the spleen) GCs. Kruskal-Wallis test followed by Dunn’s test with Benjamini-Hochberg p-value adjustment).

| Cell subsets | Comparison | Z | P.unadj | P.adj |
| --- | --- | --- | --- | --- |
| **CD20 B cells** | dLN ccRCC - dLN metPan | -0.132841 | 0.894319163 | 0.894319163 |
|  | dLN ccRCC - Spleen | -3.776181 | 0.000159252 | **0.000477755** |
|  | dLN metPan - Spleen | -3.681533 | 0.000231836 | **0.000463671** |
| **CD4 T cells** | dLN ccRCC - dLN metPan | 0.2415802 | 8.09E-01 | **8.09E-01** |
|  | dLN ccRCC - Spleen | 4.9100685 | 9.10E-07 | **2.73E-06** |
|  | dLN metPan - Spleen | 4.715811 | 2.41E-06 | **4.81E-06** |
| **CD68 macrophages** | dLN ccRCC - dLN metPan | -5.703728 | 1.17E-08 | **2.34E-08** |
|  | dLN ccRCC - Spleen | -11.031941 | 2.68E-28 | **8.04E-28** |
|  | dLN metPan - Spleen | -5.259193 | 1.45E-07 | **1.45E-07** |
| **CD8 T cells** | dLN ccRCC - dLN metPan | 0.3494185 | 7.27E-01 | 7.27E-01 |
|  | dLN ccRCC - Spleen | 4.6556622 | 3.23E-06 | **9.69E-06** |
|  | dLN metPan - Spleen | 4.3470254 | 1.38E-05 | **2.76E-05** |
| **other cells** | dLN ccRCC - dLN metPan | 0.899717 | 0.368270854 | 0.36827085 |
|  | dLN ccRCC - Spleen | -1.804539 | 0.071146862 | 0.14229372 |
|  | dLN metPan - Spleen | -2.755232 | 0.005865046 | 0.01759514 |

**Supplementary Table 2.** Multiplex immunofluorescence antibody panel, staining conditions, and corresponding figures.

| Slides | Figures | Incubation with | Antibodies | Company | Cat. No. | Working dilution [µg/mL] | Incubation time / T |
| --- | --- | --- | --- | --- | --- | --- | --- |
| ccRCC, metPan | Figure 3c; Supplementary Figure 2a and b | Primary Ab | anti-human CD20 | abcam | ab9475 |  | o.n. /+4°C |
|  |  | Primary Ab | anti-human CD3 | abcam | ab5690 | 2.0 | o.n. /+4°C |
|  |  | Secondary Ab | anti-mouse Alexa Fluor488 | Invitrogen | A32723 | 1.0 | 45 min +37°C |
|  |  | Secondary Ab | anti-rabbit Alexa Fluor555 | Invitrogen | A32733 | 0.5 | 45 min +37°C |
|  |  | Primary Ab | anti-human CA9 - Alexa Fluor647 | abcam | ab225074 | 10.0 | 45 min +37°C |
| ccRCC, metPan | Figure 3c (inset); Supplementary Figure 2b | Primary Ab | anti-human CD8 [CAL66] | abcam | ab237709 | 0.7 | o.n. /+4°C |
|  |  | Primary Ab | anti-h CD20 | abcam | ab9475 |  | o.n. /+4°C |
|  |  | Secondary Ab | anti-mouse Alexa Fluor488 | Invitrogen | A32723 | 1.0 | 45 min +37°C |
|  |  | Secondary Ab | anti-rabbit Alexa Fluor555 | Invitrogen | A32733 | 0.5 | 45 min +37°C |
| metPan, ccRCC, dLN<br>ccRCC, dLN metPan | Supplementary Figure 2c | Primary Ab | anti-human CD20 | abcam | ab9475 |  | o.n. /+4°C |
|  |  | Primary Ab | anti-CD8 [CAL66] | abcam | ab237709 | 0.7 | o.n. /+4°C |
|  |  | Primary Ab | anti-PNAd | BioLegend | 120801 | 10.0 | o.n. /+4°C |
|  |  | Secondary Ab | anti-Mouse Alexa Fluor Plus 488 | Invitrogen | A32766 | 2.0 | 45 min +37°C |
|  |  | Secondary Ab | anti-Rabbit Alexa Fluor Plus 647 | Invitrogen | A32795 | 2.0 | 45 min +37°C |
|  |  | Secondary Ab | anti-RAT Alexa Fluor594 | BioLegend | 408911 | 1.0 | 45 min +37°C |
| metPan | Supplementary Figure 15b | Primary Ab | recombinant mouse Mab-1 | Creative Biolabs | n.a. | 10.0 | o.n. /+4°C |
|  |  | Primary Ab | recombinant mouse Mab-1 GL | Creative Biolabs | n.a. | 10.0 | o.n. /+4°C |
|  |  | Primary Ab | mouse Isotype ctrl | Thermo FisherScientific | 14-4714-82 | 10.0 | o.n. /+4°C |
|  |  | Secondary Ab | anti-mouse Alexa Fluor594 | Invitrogen | A21203 | 1000.0 | 45 min +37°C |
|  |  | Secondary Ab | anti-h CA9 - Alexa Fluor647 | abcam | ab225074 | 100.0 | 45 min +37°C |
| metPan | Supplementary Figure 15e | Primary Ab | recombinant mouse Mab-1 | Creative Biolabs | n.a. | 10.0 | o.n. /+4°C |
|  |  | Primary Ab | recombinant mouse Mab-4 | Creative Biolabs | n.a. | 10.0 | o.n. /+4°C |
|  |  | Primary Ab | recombinant mouse Mab-5 | Creative Biolabs | n.a. | 10.0 | o.n. /+4°C |
|  |  | Primary Ab | mouse Isotype ctrl | Thermo FisherScientific | 14-4714-82 | 10.0 | o.n. /+4°C |
|  |  | Secondary Ab | anti-mouse Alexa Fluor594 | Invitrogen | A21203 | 1000.0 | 45 min +37°C |
|  |  | Secondary Ab | anti-human CA9 - Alexa Fluor647 | abcam | ab225074 | 100.0 | 45 min +37°C |

**Supplementary Table 3.** Schematic of CDR3 probe combination in the BaseScope assays.

| Tissue | Year | IgH CDR3 | IgL CDR3 | TCRβ CDR3 | Figures |
| --- | --- | --- | --- | --- | --- |
| ccRCC | 0 | Clone 1 VH4-34 | Clone 1 VL2-14 |  | Supplementary Figure 11a |
| dLN ccRCC | 0 | Clone 1 VH4-34 | Clone 1 VL2-14 |  | Supplementary Figure 11a |
| metPan | 16 | Clone 1 VH4-34 | Clone 1 VL2-14 |  | Figure 3b |
| dLN metPan | 16 | Clone 1 VH4-34 | Clone 1 VL2-14 |  | Figure 3b |
| ccRCC | 0 | Clone 2 VH3-72 | Clone 2 VK3-20 |  | Supplementary Figure 11d |
| dLN ccRCC | 0 | Clone 2 VH3-72 | Clone 2 VK3-20 |  | Supplementary Figure 11d |
| metPan | 16 | Clone 2 VH3-72 | Clone 2 VK3-20 |  | Supplementary Figure 11c |
| dLN metPan | 16 | Clone 2 VH3-72 | Clone 2 VK3-20 |  | Supplementary Figure 11c |
| ccRCC | 0 | Clone 1 VH4-34 |  | Clone 1 TRBV13 | Supplementary Figure 12a |
| dLN ccRCC | 0 | Clone 1 VH4-34 |  | Clone 1 TRBV13 | Supplementary Figure 12c |
| metPan | 16 | Clone 1 VH4-34 |  | Clone 1 TRBV13 | Figure 3g |
| dLN metPan | 16 | Clone 1 VH4-34 |  | Clone 1 TRBV13 | Supplementary Figure 12b |
| Spleen | 16 | Clone 1 VH4-34 |  | Clone 1 TRBV13 | Supplementary Figure 12d |
| ccRCC | 0 | Clone 1 VH4-34 |  | Clone 2 TRBV6 | Supplementary Figure 12a |
| dLN ccRCC | 0 | Clone 1 VH4-34 |  | Clone 2 TRBV6 | Supplementary Figure 12c |
| metPan | 16 | Clone 1 VH4-34 |  | Clone 2 TRBV6 | Figure 3g |
| dLN metPan | 16 | Clone 1 VH4-34 |  | Clone 2 TRBV6 | Supplementary Figure 12b |
| Spleen | 16 | Clone 1 VH4-34 |  | Clone 2 TRBV6 | Supplementary Figure 12d |
| dLN ccRCC<br>unrelated patient | n.a. | Clone 1 VH4-34 | Clone 1 VL2-14 |  | Supplementary Figure 11b |
| dLN ccRCC<br>unrelated patient |  | Clone 2 VH3-72 | Clone 2 VK3-20 |  | Supplementary Figure 11b |

**Supplementary Table 4.**
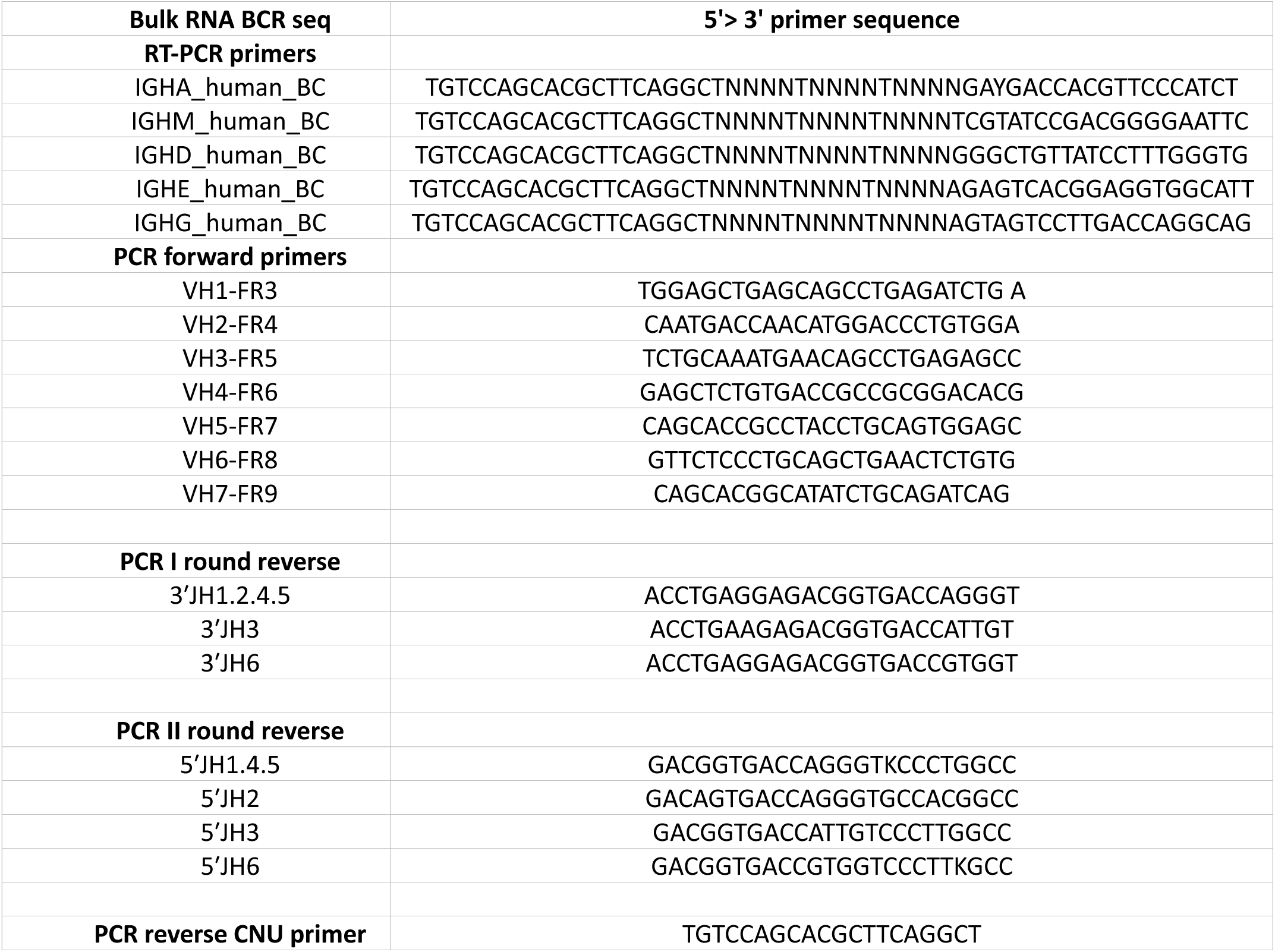
Primer sequences for bulk RNA BCR sequencing.

**Supplementary Table 5.**
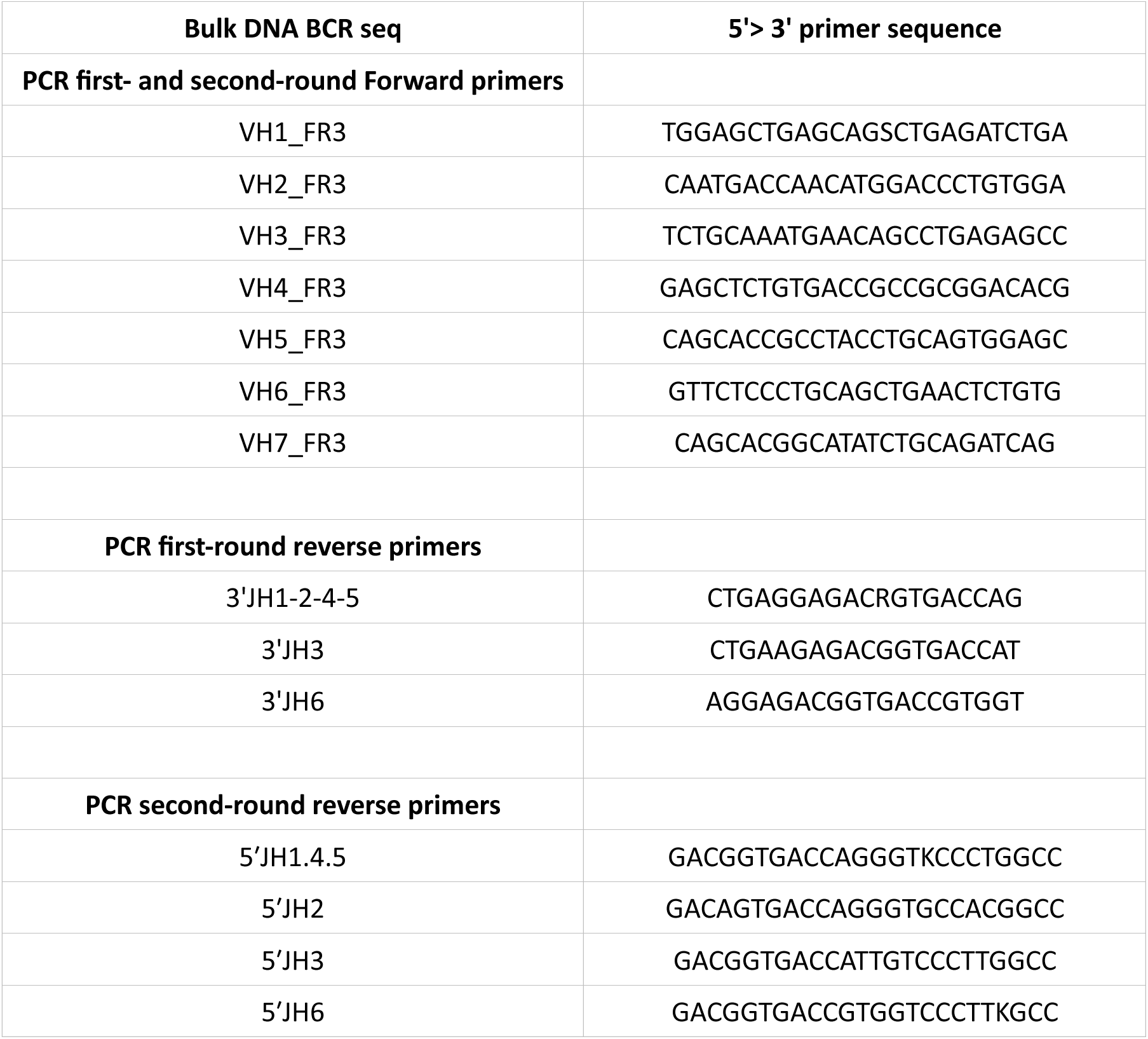
Primer sequences for bulk DNA BCR sequencing.

**Supplementary Table 6.**
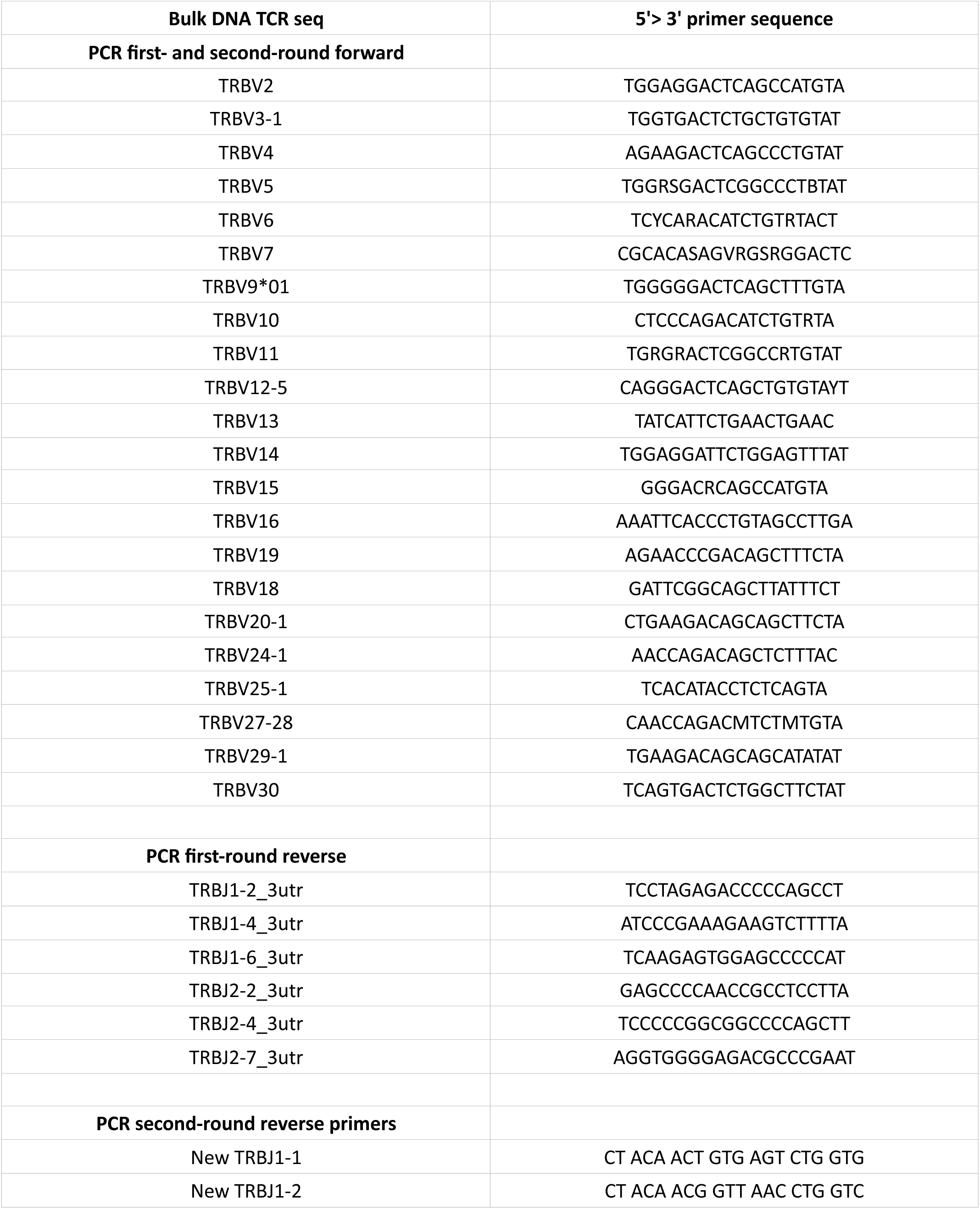
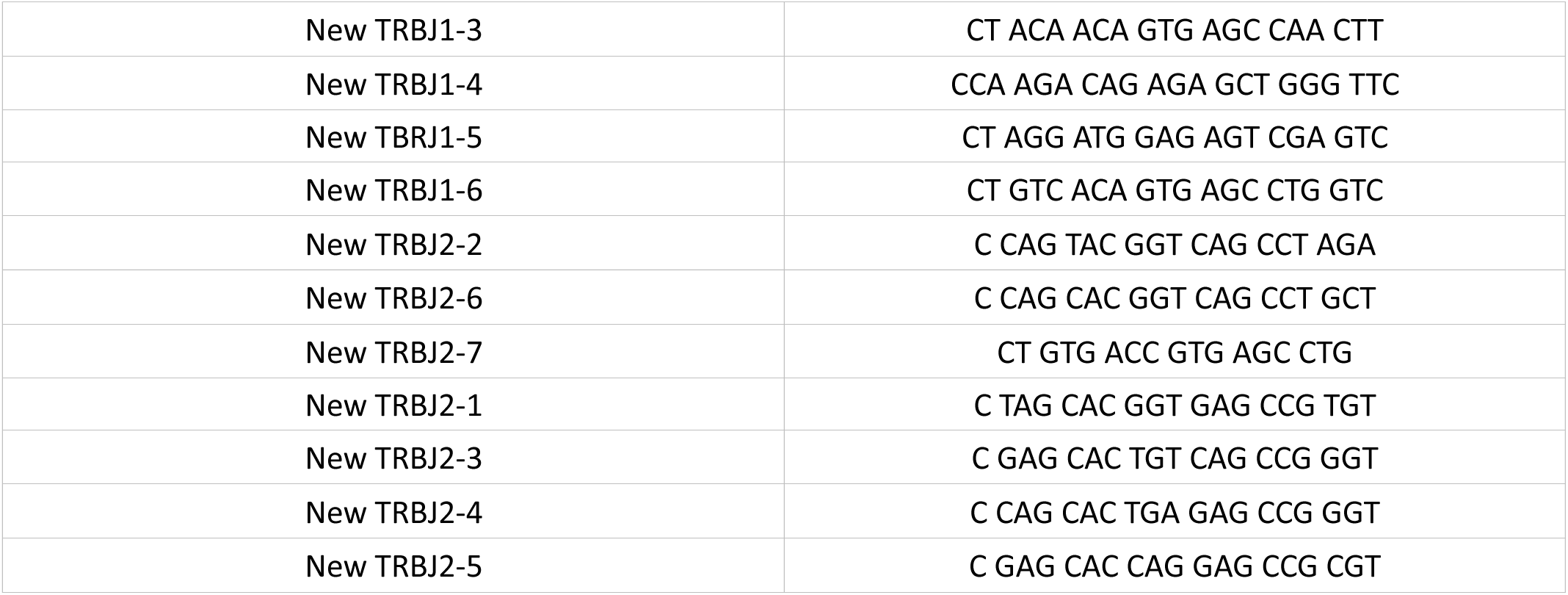
Primer sequences for bulk DNA TCR sequencing.

